# A metagenomics pipeline reveals insertion sequence-driven evolution of the microbiota

**DOI:** 10.1101/2023.10.06.561241

**Authors:** Joshua M. Kirsch, Andrew J. Hryckowian, Breck A. Duerkop

## Abstract

Insertion sequence (IS) elements are mobile genetic elements in bacterial genomes that support adaptation. We developed a database of IS elements coupled to a computational pipeline that identifies IS element insertions in the microbiota. We discovered that diverse IS elements insert into the genomes of intestinal bacteria regardless of human host lifestyle. These insertions target bacterial accessory genes that aid in their adaptation to unique environmental conditions. Using IS expansion in *Bacteroides*, we show that IS activity leads to insertion “hot spots” in accessory genes. We show that IS insertions are stable and can be transferred between humans. Extreme environmental perturbations force IS elements to fall out of the microbiota and many fail to rebound following homeostasis. Our work shows that IS elements drive bacterial genome diversification within the microbiota and establishes a framework for understanding how strain level variation within the microbiota impacts human health.

## Introduction

Bacteria rapidly evolve their genomes through the mobilization of genetic elements, including bacteriophages (phages), plasmids, DNA inversions, and transposable elements^1–3^. Genetic rearrangements have a strong impact on bacterial fitness, influencing diverse phenotypes including virulence, antibiotic resistance, interbacterial competition, and secondary metabolism. For example, pathogenicity islands can distribute virulence traits to previously non-pathogenic bacteria and DNA inversions can alter the expression of antibiotic resistance genes^4,5^. Additionally, novel genetic traits such as metabolic and virulence genes are acquired by bacteria through plasmid exchange, bacteriophage lysogeny, and conjugative transposition^6,7^. Thus, mobile elements actively drive bacterial adaptation and evolution.

Insertion sequence (IS) elements are small (∼1 kB) simple transposons that are common to bacterial genomes^8^. IS elements are capable of self-mobilization by activating an encoded transposase that recognizes sequence-specific inverted repeats at IS element termini. IS transposases are diverse^9^, and include enzymes with a DD(E/D) motif that support “copy-and-paste” and “cut-and-paste” mechanisms or those with HUH nuclease chemistry which transpose as ssDNA molecules at the replication fork^10–12^. Gene expression and genome fidelity can be modulated by IS activity in a variety of ways, including IS insertions into protein coding sequences inactivating genes, insertions into intergenic regions forming strong hybrid promoters increasing gene expression, or recombination with other IS elements resulting in large deletions^13–16^. Additionally, IS elements can rapidly expand in bacterial genomes when commensal bacteria transition to become pathogens^15,17,18^. Thus, IS elements have a profound impact on the evolution and physiological traits of bacteria^19^.

Despite a deep knowledge of fundamental IS biology from diverse bacteria, we have a rather rudimentary understanding of how these elements function in polymicrobial communities. This knowledge is critical for deciphering how bacterial communities evolve in ecosystems such as the human microbiota. A barrier to studying IS elements in complex environments result from imperfect methodologies for measuring *in situ* IS element dynamics. This stems from the poor recovery of multi-copy genes with repetitive sequences by short-read assemblers, leading to fragmented assemblies where IS elements are absent or become break-points between contigs^20^. Therefore, assembly-level analyses of IS elements from metagenomic datasets are often underpowered. Despite these limitations, previous culture-based studies show that IS elements are active in the human intestine. Sampling of *Bacteroides fragilis* isolates revealed the gain and loss of IS transposases^21^. Furthermore, IS elements were shown to rapidly expand in copy number in the genome of *Escherichia coli* during intestinal colonization of mice, driving increased virulence in a mouse model of Crohn’s disease^22,23^. Recently, longitudinal intestinal metagenomic samples from a single individual were sequenced using long DNA fragment partitioning, producing higher quality assemblies compared to short-read methodologies. This analysis revealed that *Bacteroides caccae* gained and lost numerous IS insertions over the course of sampling^20^. Finally, we previously utilized a targeted IS sequencing approach to identify insertions of the IS element IS256 in intestinal *Enterococcus faecium* populations isolated from an individual undergoing treatment with multiple antibiotics^19^. Antibiotic exposure was associated with increased abundances of IS256 insertions in genes related to antibiotic resistance and virulence, suggesting that antibiotic treatment drives IS-mediated pathoadaptation in the human intestine.

In this work, we developed an open-source IS database that greatly expands the diversity of IS elements compared to current databases. We built a computational pipeline that utilizes this database to find IS insertions in public metagenomic datasets. Our analysis reveals widespread abundance and expansion of IS insertions in the human microbiota. We show that IS insertions are transferable between individuals and are stable for years. Distinct families of IS elements are favored for insertional activity within the microbiota and these preferentially insert into specific classes of genes. Such genes are linked to distinct microbial taxa with an overrepresentation within the *Bacteroidia* and *Clostridia*. Gene classes targeted by IS elements are primarily metabolic, cell surface, and mobile genetic element genes. Using an *in vitro* ISOSDB412 (IS4351) expansion model, we confirm that identical and related accessory genes are preferentially targeted. Finally, we show that the stability of IS insertions is lost following antibiotic perturbation and diet intervention. Following these alterations, new IS abundances and insertion site locations arise. Together, this work establishes a framework for studying IS elements within the microbiota and is the first step toward understanding how IS elements contribute to the function of the microbiota impacting human health.

## Results

### ISOSDB: a comprehensive open-source IS database

To facilitate the study of IS elements in bacterial genomes and metagenomic datasets, we sought to build an updated and exhaustive IS database that can serve as an open-source tool for the scientific community. Prior to our development of this database, the main repository for IS elements was the ISFinder database^24^. While ISFinder has served as a gold-standard for the systematic naming of IS elements and their identification in single isolate genomes^25^, it is underpowered for high-throughput genomics. ISFinder lacks integration into downstream tools that can facilitate in-depth analyses of IS elements, the entirety of the database cannot be downloaded for personal use, and it relies solely on manual curation and submission of new IS elements, decreasing the speed at which new IS elements can be reported. These are drawbacks in the current genomic age, where the identification of IS elements from complex datasets such as metagenomes would greatly improve our understanding of their function.

To address the need for an unrestricted open-access IS database to support genomic research, we built the Insertion <u>S</u>equence <u>O</u>pen-<u>S</u>ource <u>D</u>ata<u>b</u>ase (ISOSDB), which consists of IS elements identified from 39,878 complete bacterial genomes and 4,497 metagenome assembled genomes (MAGs)^26^.These IS elements were identified using OASIS, a rigorously tested IS identification tool that allows for the high-throughput analysis of multiple genomes^27^. IS elements were considered valid and included in ISOSDB if: 1) there were at least two copies of the IS element in a single genome, 2) it was flanked by terminal inverted repeats (IRs), and 3) has significant nucleotide homology to an IS element in ISFinder or has a putative transposase. Redundant IS elements were deduplicated at 95% nucleotide identity. The resulting set of IS elements totaled 22,713 distinct IS elements, an almost five-fold excess to the ISFinder database (Fig. 1A). We identified transposase amino acid homologs, in addition to nucleotide homology over the whole sequence of an IS element, in the ISFinder database. 97.5% of the transposases included in ISOSDB had protein homologs in ISFinder, but only 37.9% had nucleotide homologs in ISFinder (Fig. 1B). The ISOSDB also has a wide range of transposases representing multiple IS families and contains distinct clusters of IS elements at the nucleic acid and protein level (Fig. 1C-D & Table S1). In summary, the ISOSDB is an expansive, freely available database that contains substantially improved IS diversity compared to current databases.

**Figure 1.**
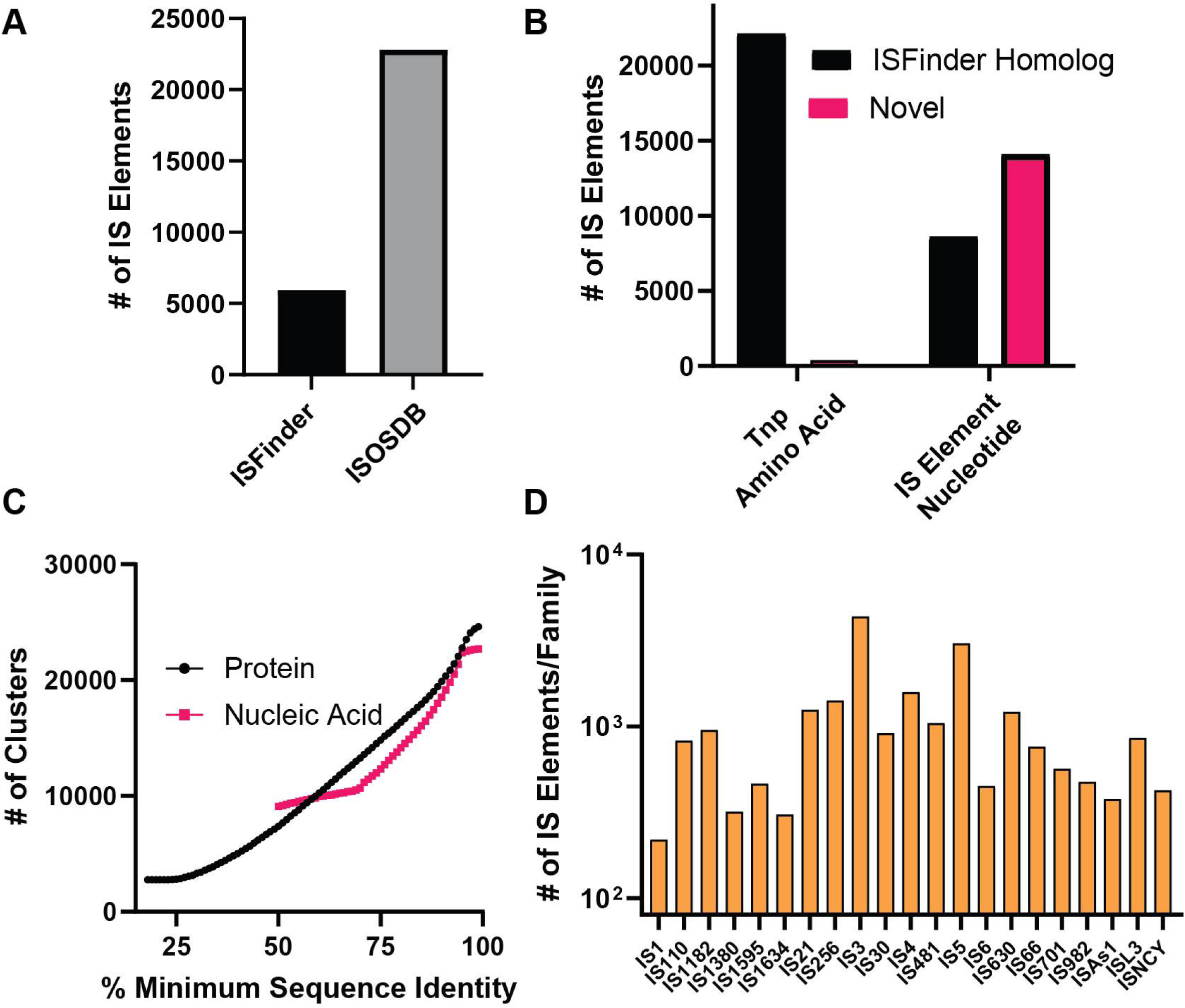
The ISOSDB: a database of diverse and complete ISs. (A) Number of unique IS sequences in ISFinder and the ISOSDB. (B) Comparison of transposase (Tnp amino acid) and nucleotide sequences of the full ISs from the ISOSDB to the ISs from ISFinder. (C) Numbers of clusters of ORFs and full nucleotide sequences in the ISOSDB for a range of minimum sequence identities. (D) Number of ISs per IS family in the ISOSDB. Only families with 100 or more IS elements are shown.

### Development of an IS detection pipeline that uses ISOSDB

IS elements drive genomic diversity in almost all bacterial species^9,28,29^. However, a systematic method to identify IS insertions in complex microbial communities has not been previously developed. This has precluded the study of IS biology in the microbiota at the population scale. To obtain a deeper understanding of IS biology in the microbiota, we developed the pseudoR pipeline that utilizes ISOSDB to identify IS insertions in previously assembled genomic sequences. pseudoR is named for its ability to find IS inactivated pseudogenes and is implemented in the R programming language. This pipeline is built for fragmented, incomplete assemblies such as metagenomes. Our approach was inspired by previous tools developed for the analysis of transposon insertions from eukaryotic^30^ and bacterial genomes^31,32^. These tools either require *a priori* knowledge of target site duplications which is often not consistent for IS insertions^33^, use read pairing to infer IS insertions which limits the data available to verify IS insertions, or require a matched reference assembly.

The backbone of the pseudoR pipeline is a split read approach. First, reads are mapped against assembled contigs and unmapped reads are binned as separate read files. These unmapped reads are aligned against a database of IS terminal ends (150 bp on either end of an IS) compiled from ISOSDB (Fig. S1). If an unmapped read has an IS termini, the termini is trimmed and the remaining read is remapped against the assembled contigs. To ensure that insertions are genuine, a minimum depth of four reads is required and these reads must include at least one left and one right termini.

To compare between samples and multiple disparate assemblies, we built two modes (multi and single). Multi-mode takes each sample and maps its reads against its own assembly. These reads are then subsequently mapped against a deduplicated gene database built from all assemblies in the dataset. Single-mode was developed to analyze time series datasets, where a single assembly can be used for multiple samples (such as one from the start of the time series).

### IS insertions are widespread in the healthy human intestinal microbiota

We utilized the pseudoR pipeline to identify IS insertions in healthy human fecal metagenomes by analyzing data from three geographically disparate studies: a survey of healthy Italian adults ranging in age from 30-105 years (ITA)^34^, healthy individuals from a Japanese colorectal cancer study (JPN)^35^, and healthy rural Madagascan adults (MDG)^36^. Each study showed evidence of widespread IS insertion heterogeneity (Fig. 2A). Some individuals had few detectable IS insertions while others had over 100 unique IS insertions (Fig. 2B). We used relative IS depth (IS depth divided by the sum of the IS depth and the depth of reads that map to the insertion site without the insertion present) as a metric to report the abundance of IS alleles. Most insertions had a relative IS depth between 10% and 100%, demonstrating that the inserted allele exists in equilibrium with the wild type allele (Fig. 2A). We next determined the bacterial taxa that underwent extensive IS insertional activity (Fig. 2C, Fig. S2A). IS insertions were most abundant in the *Bacteroidia* and *Clostridia*. These two classes make up the majority of the healthy intestinal microbiota^37^ and the genera *Bacteroides*, *Phocaeicola*, *Ruminococcus*, and *Blautia* accounted for the majority of IS element insertion diversity (Fig. S2A). While *Bacteroidia* IS insertions were present across all studies, the IS elements underlying these insertions varied between studies (Fig. 2D). Unique IS types for the *Bacteroidia* included ISOSDB412 (IS30 family) insertions for ITA individuals, ISOSDB18121 (IS1380 family) insertions for JPN individuals, and ISOSDB33 (IS1182 family) insertions for MDG individuals (Fig. 2D). Insertions in the *Bacteroidia* from JPN and ITA individuals were formed from shared IS elements, including ISOSDB4584 (ISL3 family), ISOSDB634 (IS982 family), and ISOSDB445 (IS66 family), whereas the IS diversity in the *Bacteroidia* from MDG individuals consisted of ISOSDB250 (IS66 family), ISOSDB426 (IS630 family), and ISOSDB45 (IS256 family) insertions. This suggests that regional differences contribute to IS element content or activity that drives insertion events.

**Figure 2.**
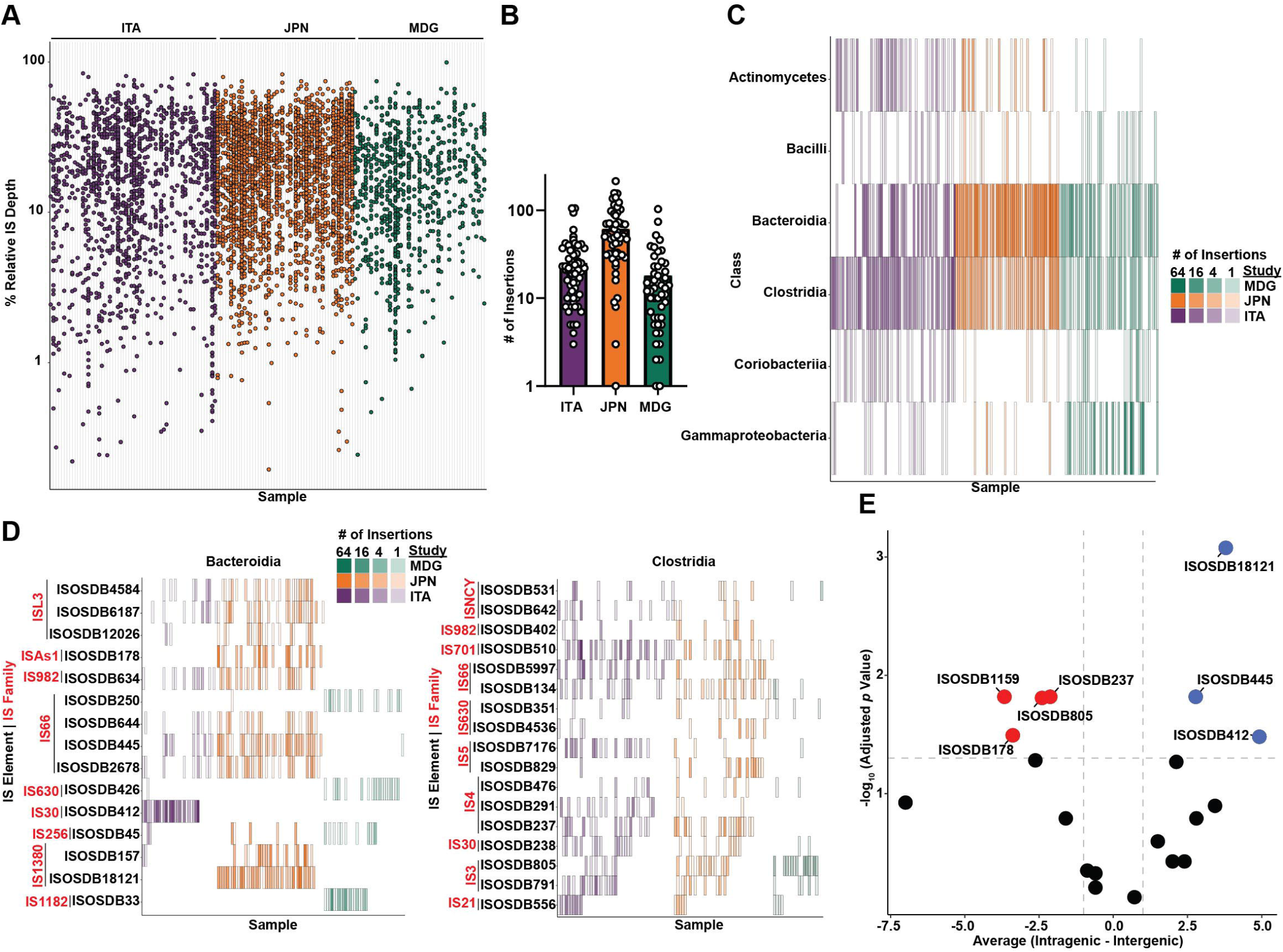
IS insertions are abundant in intestinal bacteria. (A) Abundance of ISs from ITA, JPN, and MDG individuals. Each data point represents one new insertion not found in the reference assembly. The y-axis is the relative IS depth percentage (IS depth/ (WT allele depth + IS depth)). (B) Number of insertions per sample shown in A. (C) IS insertions per bacterial class. The intensity of each bar is proportional to the number of IS insertions. (D) IS insertions in *Bacteroidia* and *Clostridia* bacteria organized by IS family. The intensity of each bar is proportional to the number of IS insertions. (E) Preferential IS insertion in either intergenic (left side, red) or intragenic (right side, blue) loci for highly abundant ISs (paired T test with FDR multiple testing correction, maximum adjusted *p* value = 0.05). The difference between intragenic and intergenic insertions individual IS elements was averaged between multiple individuals and is shown on the X axis.

IS insertion types among the *Clostridia* were found across all three geographic locations, with many more IS elements shared between JPN and ITA individuals (Fig. 2D). Interestingly, the *Clostridia* shared three IS family types (IS982, IS66, and IS30) with the *Bacteroidia*, yet the specific IS family members were different IS elements (Fig 2D). We conclude from this data that although the IS sequence space of these two classes of bacteria are unique, some IS families are shared and certain IS families may be more promiscuous.

MDG individuals, despite having similar numbers of IS insertions among their bacterial communities (Fig. 2B), had relatively low abundances of IS insertions in *Bacteroides*, *Phocaeicola, Blautia,* and *Bifidobacterium* and much higher abundance of IS elements associated with pathobionts including *Escherichia* and *Prevotella* (Fig. S2A)^38–41^. This is supported by the presence of ISOSDB45 (IS256 family) insertions that are associated with pathogenic bacteria (Fig. 2D)^19,42–44^.

IS elements can insert into coding sequences or intergenic regions of the genome which can inactivate genes or influence the expression of adjacent genes^28^. To measure IS insertions in intragenic versus intergenic sites, we analyzed the precise positions of all IS elements in ITA, JPN, and MDG individuals by comparing the number of intragenic insertions to intergenic insertions on a per individual basis. We found that specific IS elements had preference for intragenic or intergenic insertions (Fig. 2E). ISOSDB1159 (IS3 family), ISOSDB805 (IS3 family), ISOSDB178 (ISAS1 family), and ISOSDB237 (IS4 family) preferentially inserted into intergenic loci, while ISOSDB18121 (IS1380 family), ISOSDB445 (IS66 family), and ISOSDB412 (IS30 family) inserted more frequently into intragenic loci. These data indicate that some IS elements are suited to diversify genomes through mutation whereas others may preferentially insert adjacent to coding sequences to alter gene expression.

### IS elements frequently insert into accessory genes important for bacterial adaptation

Having identified numerous IS insertions within predicted coding sequences, we analyzed open reading frames (ORFs) carrying IS insertions, termed iORFs, in the ITA, JPN, and MDG individuals. A deduplicated gene database of all predicted ORFs from the metagenomes of each study were assessed for iORF’s. iORF’s with insertions from a single individual were ∼10-fold more abundant than iORFs with insertions from multiple individuals (Fig. 3A). This demonstrates that intestinal metagenomic IS insertions have a high degree of inter-individual variation. We next performed functional annotation of the iORF’s and found five broad categories that were shared between individuals and encompassed 20.7% of iORFs (Fig. 3B). Many iORFs were annotated as *susC*, *susD*, or *tonB* receptor genes in ITA and JPN individuals. These genes encode high-affinity substrate-uptake receptors for carbohydrates and cofactors^45,46^. *susC-D* homologs are frequently involved in the uptake of polysaccharides used for metabolism, with *Bacteroidia* species containing many different *susC-D* homologs^45^.

**Figure 3.**
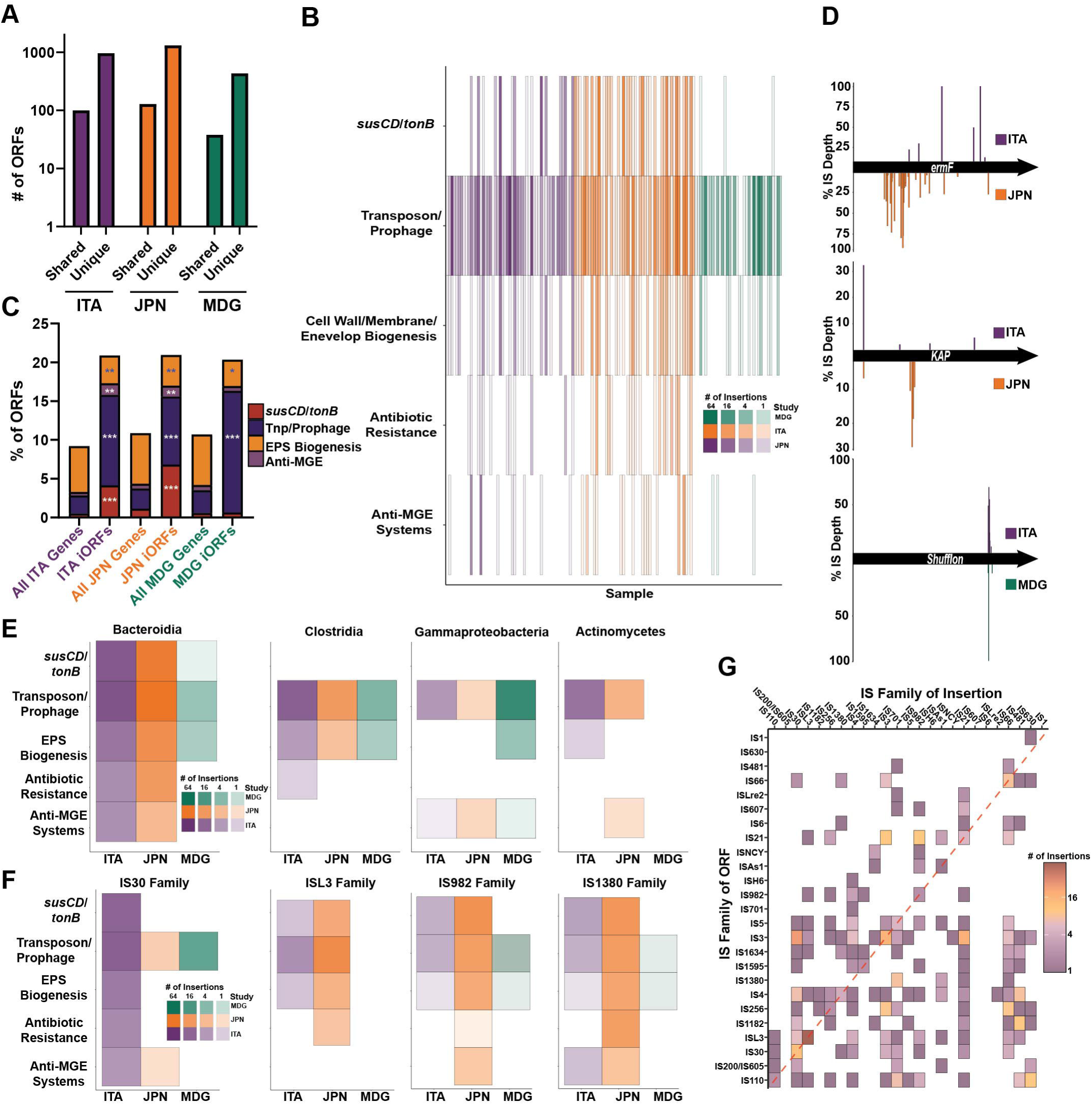
IS insertions are commonly found in bacterial accessory genes. (A) Number of shared and unique iORFs within ITA, JPN, and MDG individuals. (B) Heatmap of IS insertions in different gene functional categories. (C) Statistically significant enrichment of ISs in the functional categories from B. White asterisks represent categories where the iORFs are over-abundant compared to all genes and blue asterisks represent categories where the iORFs are under-abundant compared to all genes (Fisher’s exact test with FDR multiple comparison correction, \*\*\**p* < 10^-24^, \*\**p* < 10^-3^, \**p* < 0.05). (D) Location and relative IS depth in shared iORFs. The location of the insertion (either above or below the gene representation) is representative of the study. (E-F) Number of IS insertions in iORFs in the functional categories in B from different bacterial classes (E) and IS families (F). (G) Rate of IS insertions in transposases. Rectangles in the red dashed line are self-targets and rectangles outside of the red dashed line are non-synonymous-targets.

Mobilome genes, including transposases, integrases, prophages, and mobile element defense genes, such as restriction modification systems, contained insertions in almost every individual across all studies. Exopolysaccharide biogenesis genes, such as LPS modifying enzymes and cell wall synthesis glycosyltransferases, were consistent IS element insertion targets. Cell wall and membrane modifications are crucial for *in vivo* survival by avoiding host immunity and during competition with other bacteria^47^. Finally, antibiotic resistance genes frequently contained IS insertions, including the genes *tetQ* and *ermF* in JPN and ITA individuals. IS insertions were enriched in all functional categories except the EPS biogenesis genes, when comparing the abundance of iORFs with all predicted protein coding sequences (Fig. 3C, Fig. S2B). Together, these data show that genes involved in accessory metabolic functions, antibiotic resistance, and genomic plasticity are common IS insertion targets.

We next asked if certain iORFs were shared between individuals from the three studies. Although infrequent, a few iORFs were shared. 14% and 37% of ITA and JPN individuals, respectively, had IS insertions in the gene *ermF*, a macrolide resistance gene, and 3.2% and 9.8% of ITA and JPN individuals, respectively, had IS insertions in *KAP*, a P-loop NTPase predicted to be involved in phage defense (Fig. 3D)^48,49^. 7.9% and 6.1% of ITA and MDG individuals, respectively, shared IS insertions in *shufflon*, a gene encoding an invertase that regulates pilin phase variation (Fig. 3D)^50^. We found that iORF functional classes were conserved across diverse intestinal bacteria (Fig. 3E) yet the IS types responsible for these insertions differed among the various classes of bacteria (Fig. 3F). This demonstrates that although IS elements target similar genes within disparate bacteria, the types of IS elements that promote these diversification events are specific to certain classes of bacteria.

Multiple iORF’s were classified as transposases (Fig. 3B) and examples of IS elements inserting in other transposase genes has been previously reported^51^. In order to understand the dynamics of IS insertions in other IS elements, we classified both the transposase iORF and the IS forming the insertion by IS family (Fig. 3G). We found multiple examples of both self-targeting (an IS inserting into a closely-related IS) and orthologous-targeting (an IS element inserting into an unrelated IS). We found that IS families such as ISL3, IS4, IS110, and IS256 experienced promiscuous insertion events. These results suggest that IS self- and orthologous-targeting could be a mechanism to control IS mobilization of genetic traits.

### IS elements provide mutational diversity to Bacteroides species

Genome wide mutagenesis of *Bacteroides thetaiotaomicron* (*Bt*) identified genes that support adaptation to environmental pressures, including antibiotics, bile acids, and carbon sources^52^. We used this dataset to evaluate whether we could identify similar fitness determinants as iORFs within intestinal *Bacteroides*. We searched our iORF database for homologs to these fitness-associated ORFs from *Bt* and found multiple closely related iORFs that likely influence *Bacteroides* fitness both positively and negatively (Fig. S3A). Examples include iORFs that provide fitness advantages during growth in the presence of the antibiotics, such as doxycycline, and during hyaluronic acid and glucosamine consumption (Fig. S3A). Interestingly, we found iORFs associated with bacterial fitness during exposure to the antipsychotic medication chlorpromazine (Fig S3A) which is associated with compositional changes and antibiotic resistance of the microbiota^53–56^. Additionally, we found multiple IS insertions in homologs of the *Bt susC* gene BT1119 (Fig. S3B). Inactivation of BT1119 decreases fitness during growth on galacturonic acid^52^.

Our *in silico* analysis shows that the intestinal *Bacteroidia* experience large-scale IS mobilization into discrete groups of genes involved in carbohydrate utilization, exopolysaccharide synthesis, and mobile genetic element interactions. To test if the activation of an IS element leads to insertion into these classes of genes, we focused on ISOSDB412 (IS30 family member IS4351), an IS element is that is native to strains of *Bt* and *Bacteroides fragilis* (*Bf*)^57^. We transformed a low-copy number plasmid carrying an *ermF* cassette disrupted by ISOSDB412 into *Bt* VPI-5482 and *Bf* NCTC 9343. These strains lack native copies of ISOSDB412 which allowed us to study this IS element in the context of a genetic arrangement that we found from our analysis of the microbiota. Following growth, we isolated their genomic DNA and performed IS-Seq which enriches for ISOSDB412 amplicons to identify their insertion locations^19^. Both *Bt* and *Bf* strains carrying the ISOSDB412 plasmid obtained numerous ISOSDB412 insertions throughout their genomes compared to controls that lacked the ISOSDB412 plasmid construct (Fig. 4A, Fig. 4D). As predicted, ISOSDB412 insertions were enriched in gene categories reflected from our findings from insertions from the human microbiota (Fig. 4B, Fig. 4E). In *Bt*, ISOSDB412 insertions were enriched in *susC-D/tonB* and EPS biogenesis genes, whereas *Bf* acquired an abundance of insertions in *susC-D*/*tonB* genes despite a significantly lower number of ISOSDB412 insertion events compared to *Bt* (Fig. S4A). Additionally, insertions were more frequently found in intragenic loci compared to intergenic loci, confirming our findings from the intestinal microbiota (Fig. 4C, Fig. 4F). These results demonstrate that IS elements can rapidly diversify *Bacteroidia* genomes.

**Figure 4.**
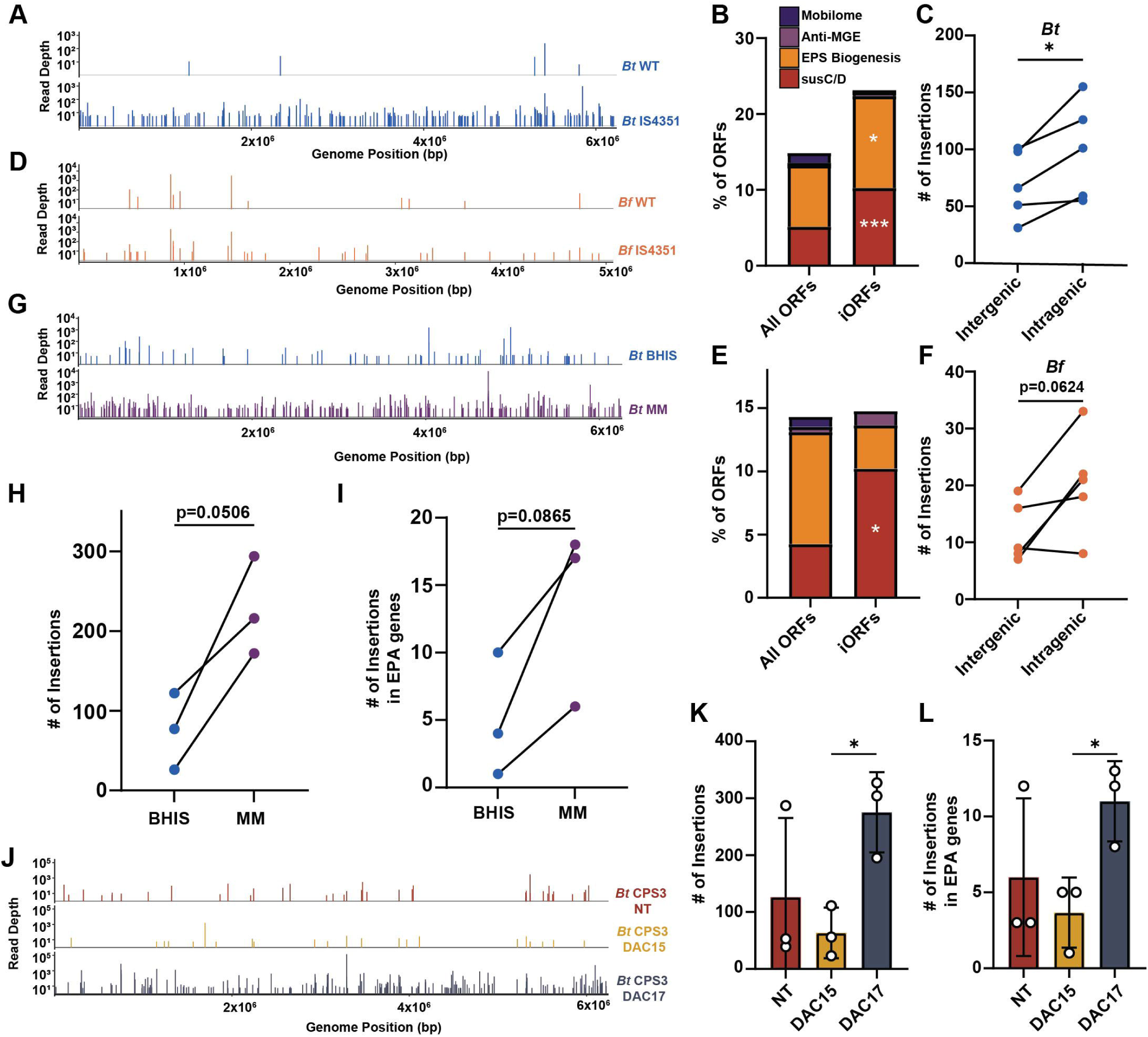
ISOSDB412 insertion in *Bacteroides* species replicates findings from intestinal metagenomic data. (A and D) Representative IS-Seq deep sequencing of ISOSDB412 insertions in *Bt* (A) and *Bf* (D). (B and E) Functional enrichment of ISOSDB412 insertions in *Bt* (B) and *Bf* (E) (Fisher’s exact test, \**p* < 0.06, \*\*\**p* < 0.001). (C and F) ISOSDB412 insertions were counted in intergenic and intragenic loci in *Bt* (C) and *Bf* (F) (paired T-test, \**p* < 0.05). (G) Representative IS-Seq of ISOSDB412 insertions of *Bt* passaged in either BHIS or MM. (H) Number of ISOSDB412 insertions in each condition in G (paired T test). (I) Number of ISOSDB412 insertions in EPS biogenesis genes in G (paired T test). (J) Representative IS-Seq of ISOSDB412 insertions of *Bt* chronically infected with DAC15 or DAC17. (K) Number of ISOSDB412 insertions in each condition in J (unpaired T test, \**p* < 0.05). (L) Number of ISOSDB412 insertions in EPS biogenesis genes in J (unpaired T test, \**p* < 0.05).

Our discovery that IS insertions target accessory genes suggests that these IS insertions inactivate genes that are currently dispensable for intestinal survival. To explore this, we measured IS insertional dynamics in a nutrient-deplete environment with glucose as the sole carbon source. We hypothesized that a lack of fiber and complex carbon sources, high levels of glucose, and single sources of iron and nitrogen, would incentivize higher rates of IS insertions to inactivate costly metabolic machinery needed for competition in complex nutritional environments. We measured ISOSDB412 insertions from three *Bt* colonies after three sequential passages in either complex media (BHIS) or minimal media (MM) (Fig. 4G, Fig. 4H, Fig. 4I). We found that colonies passaged in MM had substantially more ISOSDB412 insertions compared to BHIS-passaged colonies. Numerous insertions were found in the open reading frame BT3642, a Na^+^-dependent transporter (Fig. S4B) in MM colonies, but not in BHIS colonies. This gene has been shown to be detrimental for growth in glucose minimal media^52^.

Next, we measured ISOSDB412 abundance in *Bt* cells chronically infected with the *Crassvirales* DAC15 or DAC17 (Fig. 4J, 4K, 4L). *Crassvirales* are bacteriophages that infect *Bacteroides* species and are abundant in the human intestine^58^. *Bt* cells become chronically infected by these phages and shed them during growth (Fig. S4C). While DAC15 and DAC17 share 99.24% nucleotide identity over 98% of their genomes, DAC17 infection is associated with an increased IS insertions genome-wide (Fig. 4J, Fig. 4K) which inserted into genes involved in EPS biogenesis (Fig. 4L). Furthermore, almost all of the infected strains carried IS insertions in the CPS3 locus (Fig. S4D), which is crucial for productive phage infection and likely leads to phage resistance^59^. Together, these results show that IS diversification of the *Bt* genome is more prevalent under nutrient-limited conditions and during infection with intestinal-resident phages. This indicates that IS activity supports *Bt* adaptation when faced with fitness constraints.

### Temporal monitoring of IS activity shows that IS insertions are maintained over time

IS insertions were found in specific genetic loci coexisting with the wild type allele lacking the IS insertion, indicating that both versions of the allele are maintained in equilibrium. Previous work has shown that many phyla of intestinal bacteria harbor variable numbers of IS insertions^60^. To understand the temporal longevity of IS insertions in the microbiota, we analyzed longitudinal fecal metagenomic samples from healthy individuals^61,62^. Using a set of two individuals whose fecal samples were collected over the course of 76 and 91 weeks^61^, we measured the IS landscape of their microbiotas using pseudoR. We also analyzed longitudinal samples from an additional 10 individuals from a separate study^62^. A representative individual’s IS dynamics are in Fig. 5A-B. We used the assemblies from the first timepoint (indicated as week 0 in Fig. 5A) as the reference for the pseudoR pipeline which was then compared to all other time points to assess the maintenance of ancestral IS insertions and acquisition of new IS insertions. We found that many IS insertions are maintained for the entire time course and that some of these insertions frequently went in and out of detection (Fig. 5A). New insertions not present at week 0 arose at almost every timepoint, with a variable number of insertions per timepoint. New insertions were maintained for extended periods of time while others were only detected at the initial timepoint. These results demonstrate that IS elements are stably carried in the microbiota and are actively forming new insertions, similar to the accumulation of mutations in laboratory-evolved strains^63–65^.

**Figure 5.**
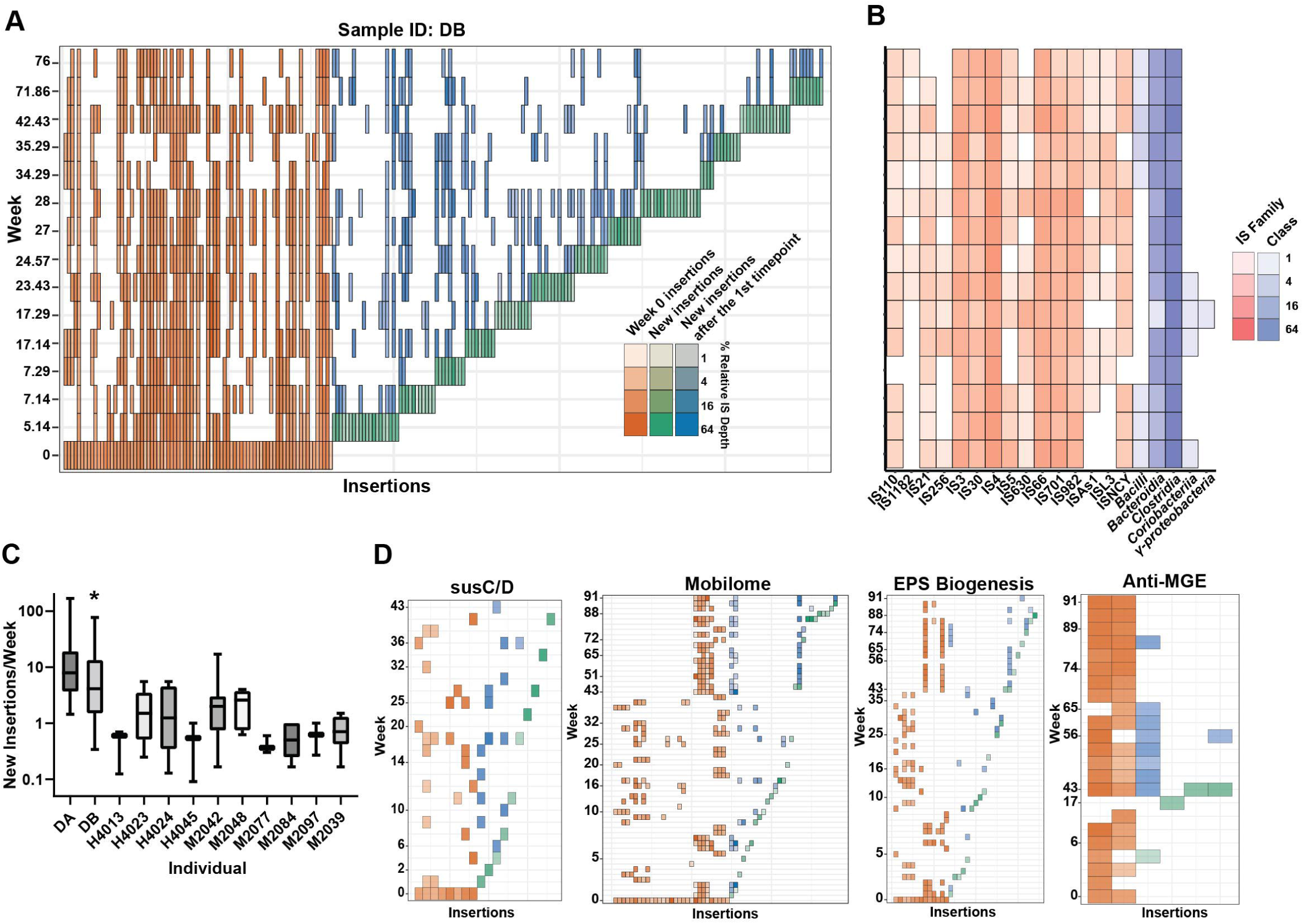
IS insertions are maintained in the intestinal microbiota for extended time periods. (A) IS insertions are depicted as rectangles, with the color intensity proportional to the relative IS depth percentage. Each insertion is unique based on insertion position and the same insertions are aligned vertically. Orange-colored IS insertions are present at the first timepoint (week 0), green IS insertions are new insertions not present at the initial timepoint, and blue IS insertions are new insertions at timepoints after their first appearance. (B) Number of IS insertions per timepoint for IS families (red) or taxonomic classes (blue). (C) Rate of new IS insertions per week. (D) Dynamics of IS insertions over time for gene functional category iORFs. Data from every individual was used for this figure. The legend for this panel is the same as in A.

Next, we analyzed whether host bacterial abundance accounts for the detection of new IS elements in all individuals (Fig. S5A). Increasing levels of host bacterial IS elements could generate higher sequencing read depth and more detection power for new IS insertions. For every insertion not present at week 0, we compared the insertion site’s read depth (a marker of host abundance) to the maximum insertion site depth of all earlier timepoints (prior to the detection of the insertion). Insertions with depths equal to or less than 200% of the previous maximum insertion site depth were considered to arise from new insertional activity (Fig. S5B). We found that the majority of new insertions could be accounted for as active IS insertions independent of bacterial abundance (Fig. S5C).

Insertions across the time series included diverse IS families and were frequently associated with the *Clostridia* and *Bacilli* (Fig. 5B). Additionally, we observed that the rate of new insertions per week for every individual ranged on average from around 1 to 10 new insertions. This value was calculated by dividing the number of new insertions per timepoint by the number of weeks between the current timepoint and the previous timepoint. We assumed a constant rate of transposition as has been used in previous studies^63,64^. Intra- and inter-personal variation in this rate could be due to both variations in selective pressure in the microbiota and technical variation, such as altered sequencing depth and library diversity. We also found that the maintenance, loss, and gain of IS elements tracked with common bacterial accessory genes that we established to be “hot spots” for IS insertions (Fig. 5C, Fig. 5D).

### IS elements are efficiently transferred and maintained within new individuals

Having discovered that individual microbiotas have unique patterns of IS insertions, we wanted to understand how the human host influences the dynamics of IS insertions within the microbiota. To test this, we used pseudoR to profile the IS insertions during fecal microbiota transplantation (FMT) where the microbiotas of FMT donors and recipients were longitudinally sampled before and after fecal transplantation^61^. We compared the IS insertional landscape in both the donors and recipients using the assembly of the donor’s transplanted microbiota as a reference. A representative recipient and donor’s IS dynamics are shown in Fig. 6A. Donor-derived communities harbored IS insertions that were stably maintained over one year in the FMT recipients (Fig. 6A). Insertions present prior to transplantation in recipients were lost, but a minority returned at later timepoints. Recipients maintained significantly less IS insertions that were present at week 0 compared to the donors (Fig. 6C), but both donor and recipients had similar rates of new insertions (Fig. 6D). To understand if IS insertional activity was similar between host and recipient, we compared the pattern of new shared insertions in donors and recipients at all timepoints post transplantation (Fig. 6B & Fig. S6). We found multiple instances of the same newly detected insertion arising in both donor and recipients at the same timepoint. These results demonstrate that IS insertions can be stably maintained in new hosts for extended periods of time and that new IS mobilization occurs at similar rates and patterns in new hosts.

**Figure 6.**
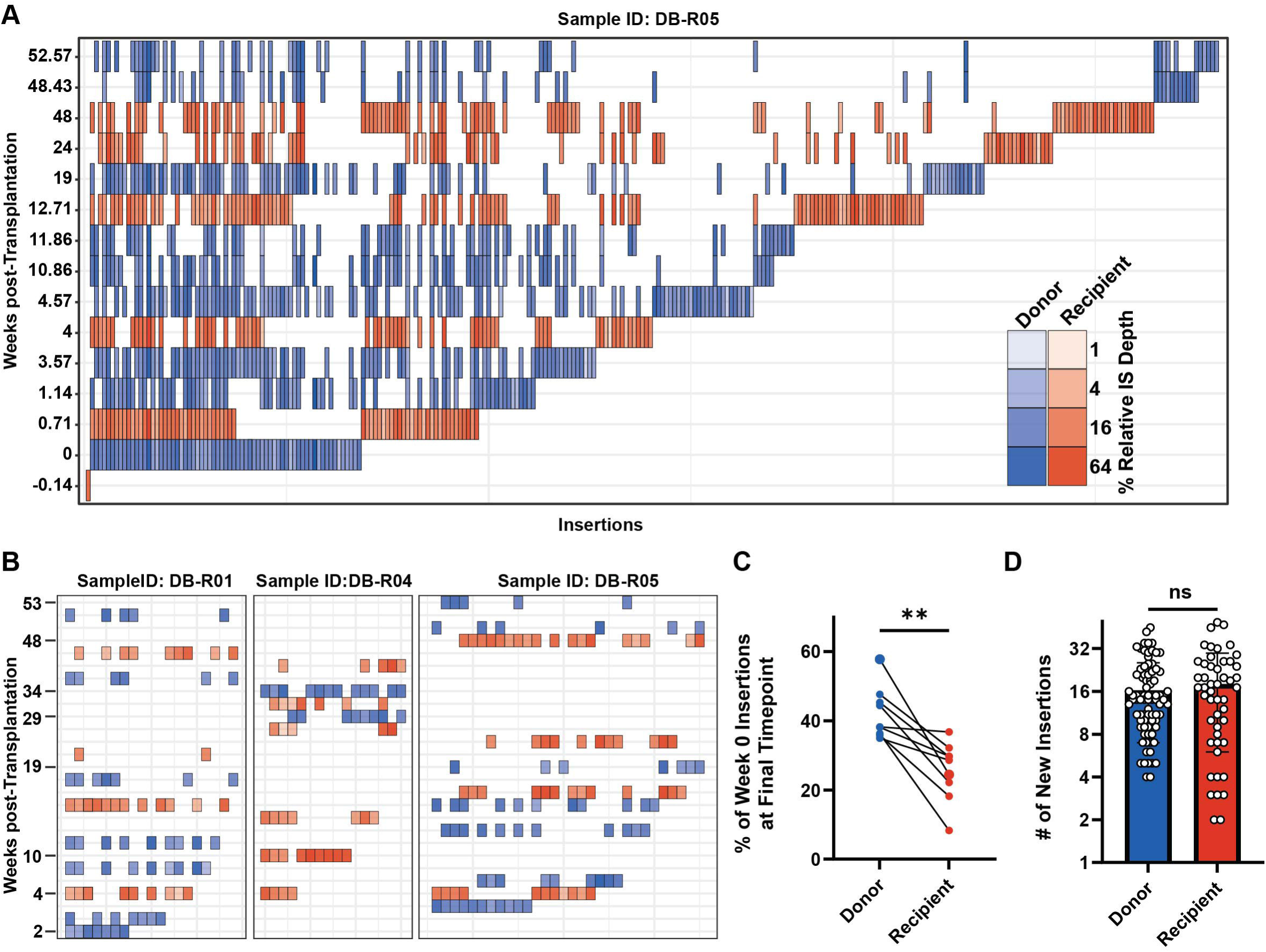
FMT transfer of IS insertions between donors and recipients. (A) Blue rectangles are IS element insertions in donor samples and red rectangles are IS insertions in recipient samples. The same insertions in donor and recipients are aligned vertically. Week 0 is the transplanted fecal material. (B) Heatmap showing new IS insertions that were detected 2 weeks after transplantation and are detected in both donor and recipient samples (see Fig. S6). (C) Loss of IS insertions detected in the transplanted fecal material (week 0) from recipients (paired T test, \*\**p* < 0.01). (D) Similar rates of new IS insertions are found in both donor and recipients (unpaired T test. Outliers removed prior to this test using the ROUT method where Q = 1%).

### Antibiotics drive widespread loss of IS insertions within the microbiota

Considering IS insertions were found to persist for prolonged periods of time within the microbiota and were portable between individuals, we wanted to know what would happen to IS diversity if a stable community was disrupted. To test this, we analyzed the IS composition before, during, and after broad spectrum antibiotic treatment^66^. The metagenome of the microbiota preceding antibiotic treatment was used as the reference for IS comparison using pseudoR. As expected, antibiotic treatment caused a substantial loss of IS diversity, which was not fully restored after treatment (Fig. 7A). The starkest example was sample ERAS10. Extensive IS diversity was present at week 0 in ERAS10 before antibiotic treatment and by at week 0.6 there were no detectable IS signatures nor did any of the original ISs return by week 26 post antibiotic therapy. To summarize these findings, we compared the rate of maintenance of pre-antibiotic treatment insertions in individuals treated with antibiotics and those who were not. Antibiotic treatment was associated with a significant decrease in the ability of the microbiota to maintain IS insertions (Fig. 7D).

**Figure 7.**
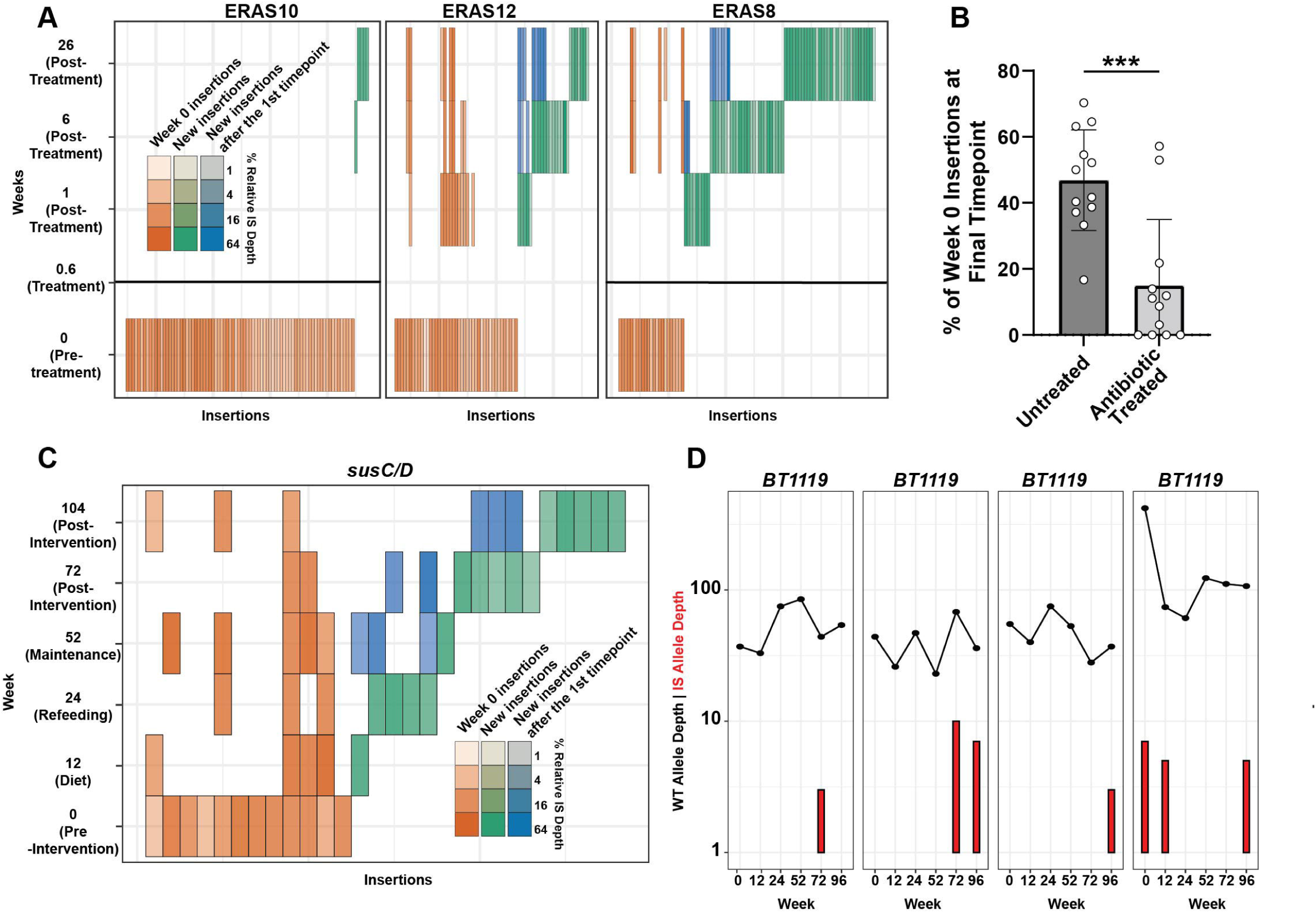
Antibiotic treatment and diet intervention alters the IS insertional landscape. (A) Heatmap showing IS insertion dynamics during antibiotic treatment. Only insertions in alleles with less than or equal to 200% of the week 0 read depth are shown in this panel. Horizontal black bars at the 0.6 Week timepoint are present in individuals who did not have an available sample for this timepoint. (B) Antibiotic treatment significantly decreases the number of IS insertions present following week 0 (unpaired T test, *** *p* < 0.005). (C) Dynamics of IS insertions in *susCD/tonB* genes during the diet timecourse. Data from every individual was used for this figure panel. (D) New *BT1119* IS insertions arise after the diet intervention has ended. Wild type allele depth is shown as a black line and IS insertion allele depth is shown as a red bar.

### IS insertion into susC-D/tonB loci is associated with diet intervention

Our analysis had identified *susC-D/tonB* genes as major targets for IS insertions, suggesting that ISs can modulate carbohydrate metabolism in the *Bacteroidia* (Fig. 3), and we showed in *Bt* that nutrient stress promotes insertions into predictable metabolic loci (Fig. 4). Based on this data, we suspected that a radical diet intervention in humans would lead to a shift in IS insertions in metabolic-associated genes such as *susC-D/tonB* loci. To test this hypothesis, we analyzed the IS insertions in the intestinal microbiota of obese individuals before, during, and after a year-long diet intervention^67^. Participants in this study received a low-calorie formula-based diet for 12 weeks, followed by a 6-week period of phasing out the formula diet for solid food without changing total caloric intake, followed by a 7-week phase where caloric intake was increased while preventing weight gain^68^. We found that ancestral *susC-D/tonB* insertions were frequently lost during the course of the diet study and new insertions in *susC-D/tonB* genes arose after the diet intervention ended (Fig. 7C). Additionally, insertions arose in the *susC- D/tonB* homolog BT1119 (Fig. 7D). Together, these results demonstrate that diet alteration is associated with perturbations in IS insertions in *susC-D/tonB* genes.

## Discussion

IS elements are fundamental to the evolution of bacterial genomes. Despite this, IS diversity and function within polymicrobial communities is understudied. Here, we built a computational pipeline for identifying IS insertions within complex metagenomic DNA sequences. This pipeline, which we call pseudoR, relies on an open-source IS element database (ISOSDB), which vastly improves nucleotide diversity compared to current state-of-the-art databases. Using pseudoR, we found that IS elements contribute to high levels of bacterial genomic diversification within the microbiota, are maintained at specific loci for extended periods of time, can be efficiently transferred between individuals, and experience fluctuations within bacterial populations following perturbation. Together, these results demonstrate that IS elements are important genetic elements that can dictate genotype diversity within the microbiota.

Using previously published metagenomic datasets, we show that the majority of open reading frames containing IS insertions (iORFs) are unique and not shared between individuals. This is similar to the observation that polymorphism diversity in the human intestinal microbiota is more likely to be unique than shared^69^. Our work expands on this idea and suggests that IS activity is a part of each person’s “fingerprint” of microbial diversity.

iORFs were frequently annotated within five broad categories: *susC-D/tonB*, mobilome genes, EPS genes, antibiotic resistance genes, and MGE resistance genes. Genes belonging to these categories are considered accessory genes and are often non-essential. Interestingly, genes regulated by DNA inversions, including phase variation, fall into similar functional categories as those targeted by IS insertions^70,71^. This suggests that insertions and inversions control gene expression of the microbiota to position bacteria for optimal adaptation. Additionally, *susC-D*/*tonB* receptors are often targets of nucleotide evolution in the human intestine, possibly to help gain new abilities based on substrate availability^21^. In support of this, a SusC epitope activates T cells to promote an anti-inflammatory IL-10 response in healthy individuals and induces a pro-inflammatory IL-17A response in people with Crohn’s disease^72^. This suggests that IS elements influence the immunomodulatory capacity of the microbiota.

The microbiome of urban individuals is less-well suited to degrade diverse polysaccharides compared to rural individuals^73^. Here, we found that urban individuals (those in the ITA and JPN cohorts) have extensive modulation of the microbiota with high rates of IS insertions in *susC-D/tonB* genes, compared to individuals living in rural settings (MDG cohort). This could partly be explained by in ITA and JPN individuals as they have a dominance of IS insertions in *Bacteroides* and *Phocaeicola* bacterial species, while MDG individuals have limited *Bacteroidia* IS insertions. However, we expect that the varied diet of MDG individuals, which includes wild meat and plants^74^, partially selects for a more diverse set of functional *susC-D/tonB* receptors compared to individuals consuming an industrialized diet. Further work to characterize IS insertions in individuals with rural lifestyles, such as the Hadza hunter gatherers^75^, will better demonstrate how IS elements impact nutrient acquisition.

iORFs were common among MGE resistance and macrolide resistance (*ermF*) genes^76^. Both of these classes of genes have been reported to be targets of IS inactivation^19,33^. Since these are heterogenous populations that co-exist between genes carrying or lacking IS insertions, we hypothesize that IS insertions in these genes maintains a low level of population-wide MGE and antibiotic resistance and reduces the burden of expressing the resistance gene. Such a “hedge-betting” strategy would be advantageous upon antibiotic or phage exposure where the population can regain resistance thorough selection of variants.

Prior to this study, it was uknown how stable IS elements were in bacterial genomes within host associated microbiotas. We found that specific IS insertions were detectable for just under two years, demonstrating that IS insertions within specific genetic loci are often not sanitized from the population. These findings are analogous to the discovery of nucleotide polymorphism conservation within the microbiota^77^. Additionally, we established that IS elements can be stably transferred between individuals. Donor-derived bacterial communities have been shown to persist in recipients following FMT^78–80^. We conclude that IS insertions are stable in host associated microbiotas and may impart a minimal fitness cost to their bacterial host.

Bacterial diversity frequently rebounds after antibiotic treatment^66,81^, but how bacterial communities assemble at the strain level is more varied and includes widespread loss of mutations and enrichment of others^82^. We found that IS insertions were readily lost after antibiotic treatment. Our results suggest that ISs can be viewed as strain-level features that are more variable than taxonomic-level diversity following perturbation.

In summary, the development of a contemporary and stringently curated IS database, combined with a streamlined computational approach to identify IS elements from metagenomic data, has revealed insights into how IS elements shape the genomes of the microbiota and sets the stage for exploring how specific IS variants in the microbiota influence human health. However, a few limitations to the tools and study exist. First, delineation of IS insertion sites between closely related strains is imperfect, thus insertions between very closely related strains may group together and fail to resolve. Work from the intestinal microbiota has found that most species are represented by one dominant strain and that these strains are stable over long periods^83^, suggesting that the insertion sites that we measured are from subpopulations originating from single strains. Second, we do not understand how IS insertions influence the fitness of the microbiota. We can begin to overcome this problem by assembling and assessing Tn-Seq libraries of diverse intestinal bacteria under conditions relevant to the intestine and in animal colonization experiments. Coupling such studies with pseudoR analyses of microbiota datasets would begin to help determine what IS insertion sites are beneficial or detrimental to commensal bacteria. Third, the pseudoR pipeline relies on assemblies containing some level of the wild type allele for comparison to an IS insertion at that same allele. This limits the detection of IS elements that assemble as part of the reference contig. Conceivably, an inverse method to find wild type alleles in IS-allele assemblies can be built and be implemented into a future release of pseudoR. Fourth, the single reference mode of pseudoR excludes assemblies from later timepoints to prevent over-assembly and focuses the analysis on the initial timepoint. This hinders our understanding of how IS elements are impacting the microbiota not present at the initial timepoints.

## Materials and Methods

### Data and Code Availability

IS-Seq data have been deposited at NCBI SRA and are publicly available as of the date of publication. Accession numbers are listed in the Key Resources Table. This paper analyzes existing, publicly available data. The accession numbers for these datasets are listed in the Key Resources Table. All original code has been deposited at GitHub and is publicly available as of the date of publication. DOIs are listed in the Key Resources Table.

### Experimental Model and Study Participant Details

*Bacteroides thetaiotaomicron* VPI-5482, *Bacteroides thetaiotaomicron* VPI-5482 CPS3, and *Bacteroides fragilis* NCTC 9343 were cultured as previously described ^84,85^. Briefly, these strains were grown in brain-heart infusion (BHI) broth (Becton Dickinson, Franklin Lakes, NJ) supplemented with 1 g/L cysteine, 5% w/v NaHCO_3_, and 5 mg/L hemin (BHIS), Varel-Bryant minimal media ^86^ supplemented with 28 mM glucose, or *Bacteroides* phage recovery medium (BPRM) ^87^ at 37° C in a Coy Type A vinyl anaerobic chamber in an atmosphere of 5% hydrogen, 20% carbon dioxide, and balanced nitrogen (Coy Lab Products, Grass Lake, MI). For growth on solid-media we used BHI agar supplemented with 10% defibrinated calf blood (Colorado Serum Company, Denver, CO). *Escherichia coli* S17-1 was grown in Lennox L broth (Fisher Scientific, Hampton, NH) with aeration at 37° C. Tetracycline was used at 2 µg/mL, gentamicin was used at 200 µg/mL, and ampicillin was used at 100 µg/mL.

### Quantification and Statistical Analysis

Details of statistical analyses for specific figures can be found in the figure legends. All statistical analyses were performed using Graphpad Prism, with the exception of Fisher’s exact test which was performed using R.

### Creation of the ISOSDB

All genomes classified as “complete” in NCBI Assembly were downloaded on 05/08/2023 using bit v1.8.57 ^88^. IS elements were identified using OASIS ^27^. This program uses annotated transposases as points for sequence extension and comparison to find complete IS elements. Identified IS elements were filtered using the following criteria: 1) multiple (minimum of 2) complete copies of the element must be present in at least one genome and 2) the elements must contain identified inverted repeat sequences. To supplement these genomes, we identified IS elements in hybrid-assembled MAGs from a large human intestinal microbiota study ^26^. These metagenomes were annotated with Prokka v1.14.6 ^89^, processed as described above, and then combined with the IS elements identified from the genomes from NCBI Assembly. Redundant IS elements in the final database were deduplicated using CD-HIT-EST^90^ with a 95% sequence cutoff and a word length of 9. IS element homologs in ISFinder were found using either blastn (complete elements) with a minimum e-value of 0.000001 and a minimum length of alignment of 224 bp or blastp (transposases) with a minimum e-value of 0.000001^91^. To further confirm that IS elements in the ISOSDB are legitimate, the transposases of IS elements that lacked nucleotide homology to IS elements in ISFinder were profiled using InterProScan ^92^ and NCBI Conserved Domain Database ^93^ using default settings. Putative transposases that lacked domains associated with transposases (such as “DDE superfamily endonuclease”, “Integrase core domain”, and “Transposase”) were removed from the final ISOSDB. The ISOSDB and pseudoR pipeline is freely available at https://github.com/joshuakirsch/pseudoR. Each IS element in this database is given a unique numeric identifier. Clustering at the amino acid and nucleotide level was performed using MMSeqs2 v15.6f452 ^94^ with coverage mode 0 and a minimum coverage of 80%.

### pseudoR pipeline

To characterize IS elements and their insertion sites in metagenomic DNA sequences we built the pseudoR pipeline. This tool relies on deeply-sequenced short read Illumina data and can be used for both metagenomic and single isolate genome data.

For this study, reads were downloaded from NCBI SRA ^95^ using fasterq-dump v2.11.0 from these studies ^34–36,61,66,67^. Study accession numbers are provided in the Key Resources Table. Duplication removal, quality trimming, and read decontamination was performed using programs from the BBTools software suite ^96^. PCR duplicated reads were removed with clumpify.sh with the flag “subs=0”. Adapter and quality trimming was performed with bbduk.sh and human, mouse, and phi29 read contamination was removed using bbsplit.sh. Cleaned and deduplicated reads were assembled using MEGAHIT v1.2.7 with the “meta-large” preset (-k- max 127 -k-min 27 -k-step 10) ^97^. All contigs used in downstream analyses were greater than 1 kB.

We built two modes for IS identification: single and multi mode. Single mode is used to identify IS insertions within the microbiota from longitudinal samples from the same individual. In this mode, only the individual’s starting timepoint assembly is used to find IS insertions. This approach reduces the chance that meaningful information is lost during clustering of multiple assemblies from multiple timepoints. The single reference mode can be used for evolution studies where a single population is compared between different treatments. Multi mode is used for comparing the insertional patterns between multiple individuals. When using the multi reference mode, reads are first mapped against the sample’s own assembly and then remapped against a combined and deduplicated ORF database built from every sample’s assembly.

Before the pipeline was initially run, a database of IS termini was built by merging the first and last 150 bp of each IS in the ISOSDB together and deduplicating these merged sequences using CD-HIT-EST ^90^ with a minimum sequence identity of 90%. The merged sequences were split into 150 bp sequences and a blastn database ^91^ was built from this deduplicated dataset.

At the beginning of the pipeline, ORFs in assemblies are predicted using pprodigal.py (a parallelizable wrapper of prodigal ^98^) and deduplicated using dedupe.sh. Reads are aligned first to the reference assembly using Bowtie2 ^99^. Any mapped read, including discordantly mapped reads, are considered positive hits. Unmapped reads are collected and aligned to the IS termini database using blastn and then filtered using a custom R script. This filtering script ensures that the IS termini is properly positioned on the read such that the remainder of the read is outside of the IS element. The IS termini are removed and the trimmed reads are re-mapped to the sample’s assembly as unpaired reads using Bowtie2. Following this, the original mapped reads and IS-termini trimmed reads are re-mapped against the deduplicated gene database in nucleotide format.

Following read mapping completion in both single and multi mode, a custom R script is used to identify DNA insertion sites with read mapping from both the left and right terminus of the IS element at most 20 bp away from one another and this script compiles read abundances for these regions. Mosdepth v0.3.3^100^ is used to determine the sequencing depth of the uninserted allele for the loci described above. Seqkit v2.2.0^101^, samtools v1.6^102^, R, ggplot2, tidyr, and MetBrewer are used for various computational tasks. Scripts to reproduce all data analysis presented here are available in the GitHub repository.

### iORFs assoated with *B. thetaiotaomicron*

*B. thetaiotaomicron* VPI-5482 coding sequences were downloaded from fit.genomics.lbl.gov. Homologs to iORFs were found using blastp with a maximum e-value of 0.00005 and a minimum identity of 75%. Hits to multiple related fitness determinants were filtered to include the representative with the strongest fitness value.

#### Functional and taxonomic classification of contigs and ORFs

Metagenomic sequences with IS insertions were functionally annotated using eggNOG mapper v2^103^. Orthologous groups were cross-validated using the COG database^104^. ORFs were classified as *susCD/tonB* if they contained a PFAM which contained the phrases “SusC”, “SusD”, or “tonB”. Anti-MGE ORFs were classified as such if they classified as COG5340, COG2189, COG0286, COG0732, COG0827, or COG4217. Exopolysaccharide biogenesis ORFs and mobile element ORFs were classified as belonging to COG category “M” or ‘X”, respectively. Antibiotic resistance genes were identified using CARD RGI 6.0.0^105^. ORFs and contigs containing IS insertions were classified using Kraken2 v2.0.7^106^.

### Bacteria

*Bacteroides thetaiotaomicron* VPI-5482, *Bacteroides thetaiotaomicron* VPI-5482 CPS3^85^, and *Bacteroides fragilis* NCTC 9343 were cultured as previously described^84^. Briefly, these strains were grown in brain-heart infusion (BHI) broth (Becton Dickinson, Franklin Lakes, NJ) supplemented with 1 g/L cysteine, 5% w/v NaHCO_3_, and 5 mg/L hemin (BHIS), Varel-Bryant minimal media^86^ supplemented with 28 mM glucose, or *Bacteroides* phage recovery medium (BPRM)^87^ at 37° C in a Coy Type A vinyl anaerobic chamber in an atmosphere of 5% hydrogen, 20% carbon dioxide, and balanced nitrogen (Coy Lab Products, Grass Lake, MI). For growth on solid-media we used BHI agar supplemented with 10% defibrinated calf blood (Colorado Serum Company, Denver, CO). *Escherichia coli* S17-1 was grown in Lennox L broth (Fisher Scientific, Hampton, NH) with aeration at 37° C. Tetracycline was used at 2 µg/mL, gentamicin was used at 200 µg/mL, and ampicillin was used at 100 µg/mL.

### DNA extraction and IS-Seq

Genomic DNA was extracted from 1.5 mL cultures of *Bacteroides* strains using the Zymobionics DNA Miniprep kit (Zymo Research, Irvine, CA) and used as input for IS-Seq as described previously ^19^. Briefly, NGS libraries were produced using the Illumina DNA Prep Kit (Illumina, San Diego, CA) and amplified using ISOSDB412 IS-Seq Step1 and p7 primers for 13 cycles using Q5 Master Mix (New England Biolabs, Ipswich, MA). The products of this reaction were amplified with IS-Seq Step2 and p7 primers for 9 cycles using Q5 Master Mix and sequenced on a Novaseq 6000 at Novogene (Sacramento, CA). R1 reads were binned and trimmed of the ISOSDB412 terminus and adapters using cutadapt v1.1.18 ^107^. To remove PCR duplicates, R2 pairs of R1 reads that contained the ISOSDB412 terminus were binned and deduplicated using dedupe.sh from the BBTools suite with the flag minidentity=100.

Deduplication of R2 reads ensures that each amplified read pair is a unique molecule, while maintaining read depth information of the R1 (IS-amplified) read. This set of deduplicated R2 reads was re-paired to trimmed R1 reads with reformat.sh from the BBTools suite and these pairs were aligned to the host genome using Bowtie2. R1 read depth at each genomic position was calculated using bedtools v2.30.0 ^108^ IS-seq reads can be found at the European Nucleotide Archive under study accession number PRJEB66483.

### DNA manipulation and cloning

Primers and plasmids used in this study are listed in the Key Resources Table. All plasmid constructs were made using Gibson assembly master mix (New England Biolabs, Ipswich, MA). Assembled constructs were electroporated into electrocompetent *E. coli* S17-1. These cells were prepared by washing cell pellets three times in ice-cold electroporation buffer (0.5 M sucrose, 10% glycerol). The resulting transformants were mated with *B. thetaiotaomicron* or *B. fragilis* by mixing 500 µL of stationary phase *E. coli* S17-1 and 500 µL of stationary phase *Bacteroides* on BHI blood agar overnight at 37° C under aerobic conditions. Transconjugants were selected by scraping the bacterial lawns into BHI and plating serial dilutions on BHI blood agar supplemented with tetracycline and gentamicin, and incubating at 37°C anaerobically for 48 hours. The ISOSDB412 (IS4351) DNA sequence construct was purchased from Twist Biosciences (San Francisco, CA). pB006 (Addgene plasmid #182320), a gift from Dr. Lei Dai, was used as a shuttle vector and purchased from Addgene (Watertown, MA)^109^. The *tetM* cassette was cloned from pCIE-*tetM*^110^ and the *ermF* cassette was cloned from pG10K, a gift from Dr. Janina Lewis (Addgene plasmid #191377) and purchased from Addgene^111^.

### *B. thetaiotaomicron* passaging experiment

*B. thetaiotaomicron* was grown overnight in BHIS and then subcultured 1:100 into either Varel-Bryant minimal media or BHIS containing antibiotic and grown overnight. The next day, bacteria cultured in BHIS were subcultured 1:100 into BHIS and bacteria cultured in Varel-Bryant minimal media cultured were subcultured 1:100 into Varel-Bryant minimal media with antibiotic. After overnight growth, this passaging was repeated once more. Genomic DNA was isolated and the final passage was sequenced as described above.

### B. thetaiotaomicron phage infection

*B. thetaiotamicron* CPS3 (a mutant of *B. thetaiotamicron* VPI-5482 expressing only the CPS3 capsule)^85^ was transformed with pB6T-ermF-IS4351 and grown overnight in *Bacteroides* phage recovery medium (BPRM) with antibiotic. Overnight cultures were diluted to OD_600_ of 1.0 in phage buffer^59^ and was plated on BPRM agar containing tetracycline and 1×10^8^ PFU/mL of either DAC15 or DAC17. Dilutions of resuspended overnight culture were also plated on BPRM containing tetracycline. Plated cultures on DAC15 agar media were incubated at 37° C for 48 hours and individual colonies were picked and patched onto fresh BPRM plates with DAC15. This process was repeated. Cultures plated on DAC17 agar and agar without phage were similarly passaged. Passaged colonies were cultured into BPRM broth without phage and incubated for 48 hours at 37° C. Genomic DNA was isolated and sequenced as described above.

### Enumeration of infectious phage particles

90 µL of supernatant from liquid cultures of passaged colonies were mixed with 10 µL chloroform (Fisher Scientific), centrifuged for 1 minute at 21,000 RCF at 25° C, and the supernatant was collected. Dilutions of this supernatant were made in phage buffer and 5 µL spots of the dilutions were plated on a BPRM plate containing *B. thetaiotamicron* CPS3 embedded in 0.35% BPRM top agar, and incubated at 37° C overnight.

## Supporting information

Supplementary Key Resources Table

## Acknowledgements

We would like to thank the members of the Duerkop lab, Dr. A. Murat Eren, and Dr. Alejandro Reyes Muñoz for providing useful insights related to this project. This work was supported by the National Institutes of Health grants R01AI141479 (B.A.D.), T32AI052066 (J.M.K), F31AI169976 (J.M.K.), and R35GM150996 (A.J.H). The funders had no role in the study design, data collection and analysis, decision to publish, or preparation of the manuscript.

## Author Contributions

Conceptualization, J.M.K., B.A.D.; Methodology, J.M.K., B.A.D.; Software, J.M.K.; Validation, J.M.K.; Formal Analysis, J.M.K.; Investigation, J.M.K.; Resources, B.A.D., A.J.H.; Data Curation, J.M.K.; Writing – Original Draft, J.M.K., A.J.H., B.A.D.; Writing – Review & Editing, J.M.K., B.A.D.; Visualization, J.M.K., B.A.D.; Supervision, B.A.D.; Project Administration, B.A.D.; Funding Acquisition, J.M.K., A.J.H., B.A.D.

## Supplemental Tables

**Table S1. IS element families represented in ISOSDB, Related to Figure 1>**

