## Supplementary Key Resources Table for "A metagenomics pipeline reveals insertion sequence-driven evolution of the microbiota"

| **Bacterial and virus strains** | | |
| --- | --- | --- |
| *Escherichia coli* S17-1 | Lora V. Hooper | ATCC 47055 |
| *Bacteroides thetaiotaomicron* VPI-5482 | Lora V. Hooper | ATCC 29148 |
| *Bacteroides thetaiotaomicron* VPI-5482 CPS3 | (1) (Hickey, 2015) | N/A |
| *Bacteroides fragilis* NCTC 9343 | Lora V. Hooper | ATCC 25285 |
| *Crassvirales* DAC15 | (2) (Hryckowian, 2020) | N/A |
| *Crassvirales* DAC17 | (2) (Hryckowian, 2020) | N/A |
| **Chemicals, peptides, and recombinant proteins** | | |
| Brain heart infusion media | Becton Dickinson | 237200 |
| Ampicillin | Research Products International | A40040-25 |
| Tetracycline | Research Products International | T17000-25.0 |
| Erythromycin | Research Products International | E57000-25.0 |
| Glucose | Thermo-Fisher | D16-500 |
| Cysteine | MP Biomedicals | 101444 |
| LB broth, Lennox | Thermo-Fisher | BP1427-500 |
| LB agar, Lennox | Thermo-Fisher | BP9745-500 |
| Varel-Bryant minimal media | (3) (Varel, 1974) | N/A |
| Bacteroides phage recovery media | (4) (Tartera, 1992) | N/A |
| Bacteroides phage buffer | (2) (Hryckowian, 2020) | N/A |
| **Critical commercial assays** | | |
| Illumina DNA Prep kit | Illumina | 20060060 |
| ZymoBIONICS DNA Miniprep Kit | Zymo Research | D4300 |
| **Deposited data** | | |
| Shotgun gut metagenomics of Italian individuals | (5) (Rampelli, 2020) | PRJNA553191 |
| Shotgun gut metagenomics of Japanese individuals | (6) (Yachida, 2019) | PRJDB4176 |
| Shotgun gut metagenomics of Madagascarian individuals | (7) (Pasolli, 2019) | PRJNA485056 |
| Longitudinal shotgun gut metagenomics of American individuals | (8) (Lloyd-Price, 2019) | PRJNA398089 |
| Longitudinal shotgun gut metagenomics of American individuals undergoing FMT | (9) (Watson, 2023) | PRJNA701961 |
| Longitudinal shotgun gut metagenomics of Danish individuals undergoing antibiotic treatment | (10) (Palleja, 2018) | PRJEB20800 |
| Longitudinal shotgun gut metagenomics of German individuals undergoing diet intervention | (11) (Louis, 2016) | PRJNA290729 |
| Human reference genome NCBI build 37, GRCh37 | Genome Reference Consortium | http://www.ncbi.nlm.nih.gov/projects/genome/assembly/grc/human/ |
| Mouse reference genome NCBI build 39, GRCm39 | Genome Reference Consortium | https://www.ncbi.nlm.nih.gov/grc/mouse |
| IS-Seq of *Bacteroides* strains | This study | PRJEB66483 |
| **Oligonucleotides** | | |
| pB6_tetM_fwd  aataacttagaaacaataggccacatgc | Integrated DNA Technologies (IDT) | pB6T Gibson primer |
| pB6_tetM_rev  taattttcattataacctctccttaatttattgc | IDT | pB6T Gibson primer |
| tetM_fwd  agaggttataatgaaaattattaatattggagttttagc | IDT | pB6T Gibson primer |
| tetM_rev  cctattgtttctaagttattttattgaacatatatcgtac | IDT | pB6T Gibson primer |
| ErmF_fwd  atcgataagcttgatatcgagctcatctgcaacttttttttc | IDT | pB6T-ermF primer |
| ErmF_rev  acggctgacatgggaattccctacgaaggatgaaatttttc | IDT | pB6T-ermF primer |
| pB6_ermF_fwd  ggaattcccatgtcagcc | IDT | pB6T-ermF primer |
| pB6_ermF_rev  tcgatatcaagcttatcgatac | IDT | pB6T-ermF primer |
| Upstream Frag_fwd  aaactcccaaagtgtggggactaaactcccaaagtg | IDT | Gibson primer for pB6T-ermF-IS4351 |
| Upstream Frag_rev  aagttgaactcaagccataaggaatatttgacaccac | IDT | Gibson primer for pB6T-ermF-IS4351 |
| IS4351_fwd  ggtgtcaaatattccttatggcttgagttcaacttataaatgc | IDT | Gibson primer for pB6T-ermF-IS4351 |
| IS4351_rev  aaaatatcggaagtaatgccagctgaattcaacttgcaaatg | IDT | Gibson primer for pB6T-ermF-IS4351 |
| aagttgaattcagctggcattacttccgatattttcaaaatc | IDT | Gibson primer for pB6T-ermF-IS4351 |
| Downstream Frag_rev  gcccggctgacgccgttggatacaccaaggaaagtc | IDT | Gibson primer for pB6T-ermF-IS4351 |
| pB6T_fwd  actttccttggtgtatccaacggcgtcagccgggca | IDT | Gibson primer for pB6T-ermF-IS4351 |
| pB6T_rev  actttgggagtttagtccccacactttgggagtttaaaagtaaattagtccccacac | IDT | Gibson primer for pB6T-ermF-IS4351 |
| **Recombinant DNA** | | |
| Plasmid: pB006 | Addgene | 182320 |
| Plasmid: pG10K | Addgene | 191377 |
| Plasmid: pCIE-tetM | Gary M. Dunny | N/A |
| Plasmid: pB6T | This study | N/A |
| Plasmid: pB6T-ermF | This study | N/A |
| Plasmid: pB6T-ermF-IS4351 | This study | N/A |
| Q5 Master Mix | New England Biolabs | M0492S |
| **Software and algorithms** | | |
| pseudoR | This study | https://github.com/joshuakirsch/pseudoR |
| bit (v1.8.57) | (12) (Lee, 2022) | https://github.com/AstrobioMike/bit |
| OASIS | (13) (Robinson, 2012) | https://github.com/dgrtwo/OASIS |
| Prokka (v1.14.6) | (14) (Seeman, 2014) | https://github.com/tseemann/prokka |
| CD-HIT-EST (v4.8.1) | (15) (Li, 2006) | https://github.com/weizhongli/cdhit |
| blastp (v2.14.1+) | (16) (Altschul, 1990) | https://www.ncbi.nlm.nih.gov/books/NBK279690/ |
| fasterq-dump (v2.11.0) | NCBI, SRA Toolkit | https://github.com/ncbi/sra-tools |
| BBTools | (17) (Bushnell, 2023) | https://jgi.doe.gov/data-and-tools/software-tools/bbtools/ |
| MEGAHIT (v1.2.7) | (18) (Li, 2015) | https://github.com/voutcn/megahit |
| prodigal | (19) (Hyatt, 2010) | https://github.com/hyattpd/Prodigal |
| Bowtie2 (v2.4.5) | (20) (Langmead, 2012) | https://bowtie-bio.sourceforge.net/bowtie2/index.shtml |
| Mosdepth (v0.3.3) | (21) (Pedersen, 2018) | https://github.com/brentp/mosdepth |
| Seqkit (v2.2.0) | (22) (Shen, 2016) | https://bioinf.shenwei.me/seqkit/ |
| R (v4.3.0) | R Project for Statistical Computing | https://www.r-project.org/ |
| ggplot2 (v3.4.2) | Hadley Wickham | https://ggplot2.tidyverse.org/ |
| tidyr (v1.3.0) | Hadley Wickham, Davis Vaughan, Maximilian Girlich | https://tidyr.tidyverse.org/ |
| MetBrewer (v .2.0) | Blake R. Mills | https://github.com/BlakeRMills/MetBrewer |
| eggNOG (v2) | (23) (Cantalapiedra, 2021) | http://eggnog-mapper.embl.de/ |
| CARD RGI (v6.0.0) | (24) (Alcock, 2020) | https://card.mcmaster.ca/analyze/rgi |
| Kraken2 (v2.0.7) | (25) (Wood, 2019) | https://github.com/DerrickWood/kraken2 |
| InterProScan (v5.66-98.0) | (26) (Zdobnov, 2001) | https://github.com/ebi-pf-team/interproscan |
| NCBI CDD | (27) (Marchler-Bauer, 2011) | https://www.ncbi.nlm.nih.gov/Structure/cdd/cdd.shtml |
| MMSeqs2 (v15.6f452) | (28) (Steinegger, 2017) | https://github.com/soedinglab/MMseqs2 |
| Samtools (v1.6) | (29) (Li, 2009) | http://www.htslib.org/ |
| COG Database | (30) (Tatusov, 2000) | https://www.ncbi.nlm.nih.gov/research/cog-project/ |
