## Supplementary Figures for "A metagenomics pipeline reveals insertion sequence-driven evolution of the microbiota"

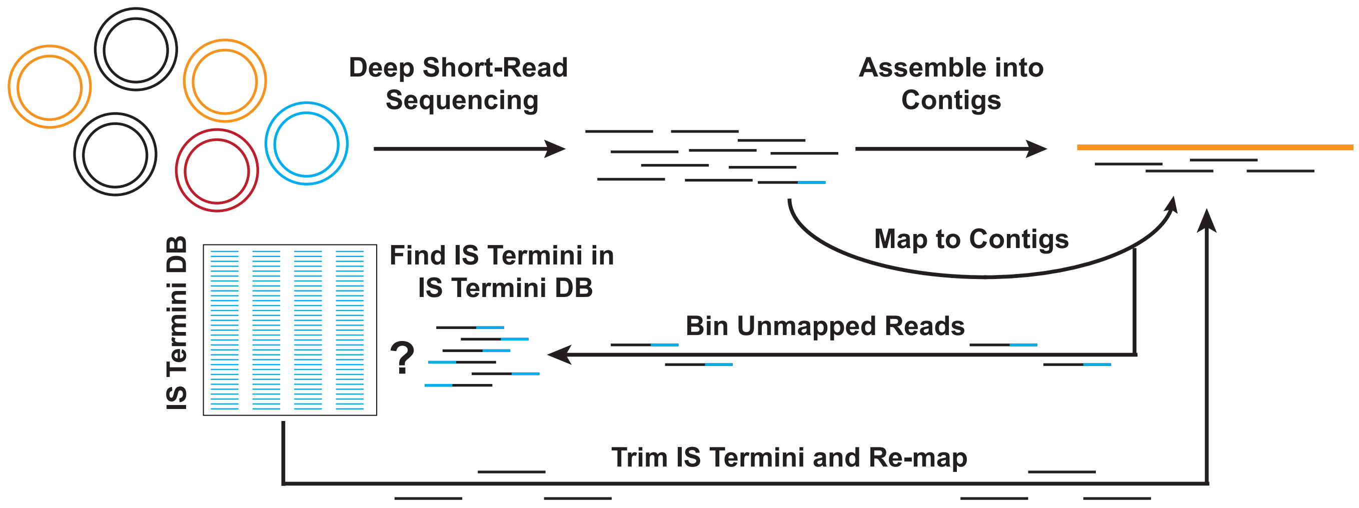


**Figure S1. Schematic of the pseudoR computational pipeline, related to STAR Methods**. Bacterial genomes are sequenced using Illumina short-read DNA sequencing. These reads are assembled into contigs and the reads (black rectangles) are mapped against the assembly contigs. Unmapped reads (black and blue lines) are compared against the ISOSDB containing IS element termini. IS element termini are trimmed from the reads and the reads are then are re-mapped to the assembly contigs.

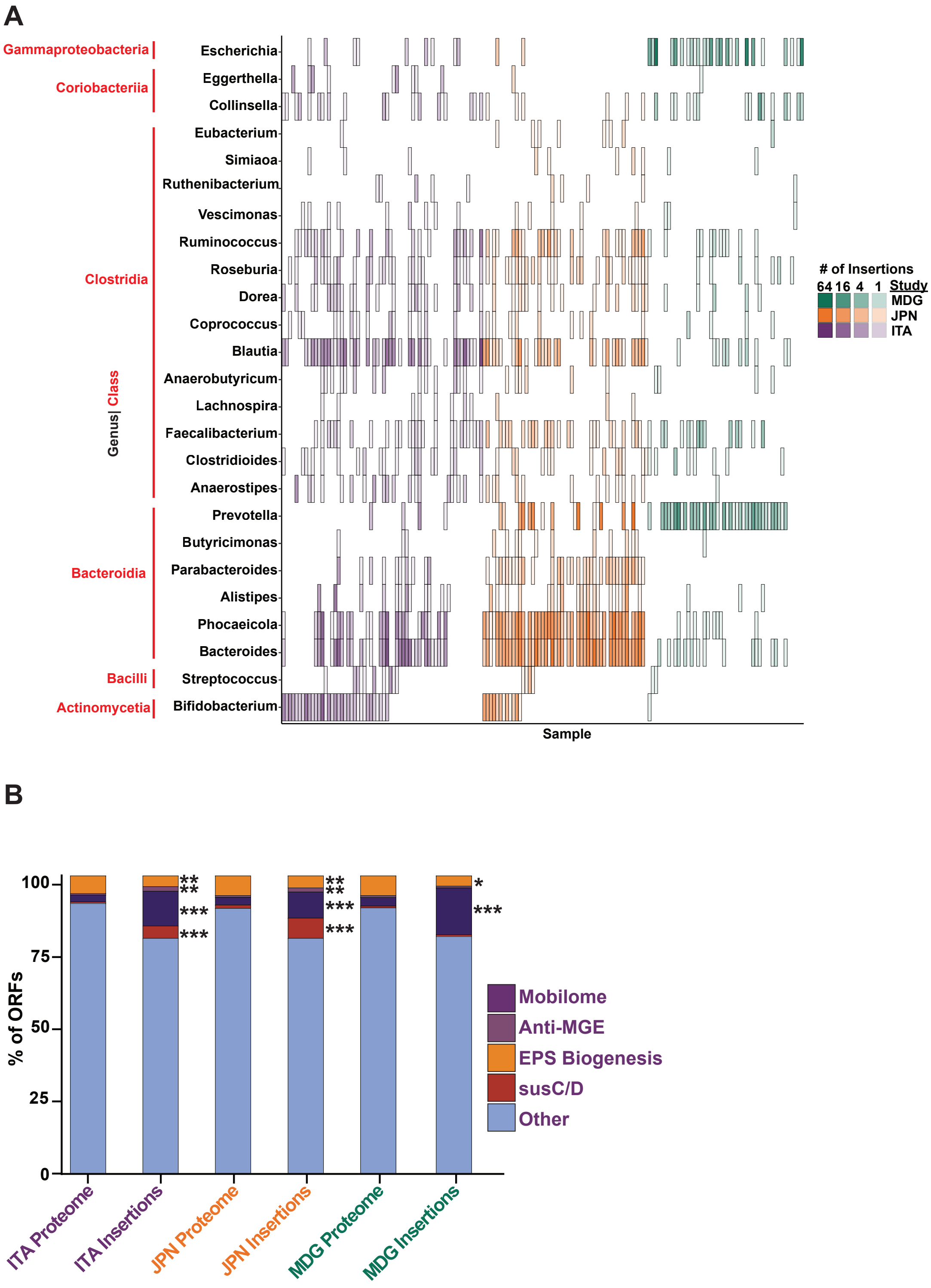


**Figure S2. IS element insertions in diverse genera and functional classes, related to Figures 2-3.**

**(A)** Abundance of IS element insertions for intestinal bacteria at the genera level. **(B)** Replotting of the data in Fig. 3C to include the "other" category which encompasses all other genes containing IS element insertions that were not the focus of this study (Fisher's exact test with FDR multiple comparison correction, ***$p < 10^{-24}$, **$p < 10^{-3}$, *$p < 0.05$).

**A**

| *Bt* Gene | % Identity | Condition | Fitness | # of Insertions |
|---|---|---|---|---|
| BT3152 | 88.126 | Doxycycline hyclate | +++ | 1 |
| BT0598 | 94.172 | Hyaluronic acid | ++ | 1 |
| BT4656 | 83.984 | Glucosamine Hydrochloride | ++ | 1 |
| BT1920 | 81.481 | Vancomycin | ++ | 1 |
| BT0602 | 81.373 | Hyaluronic acid | + | 1 |
| BT3003 | 100 | Doxycycline hyclate | + | 1 |
| BT4609 | 78.056 | Chlorpromazine hydrochloride | + | 1 |
| BT4213 | 82.51 | Chlorpromazine hydrochloride | – | 1 |
| BT2400 | 91.534 | Dimetridazole | – | 1 |
| BT1119 | 99.406 | Galacturonic Acid monohydrate | – – | 5 |
| BT4409 | 79.603 | Dimetridazole | – – | 1 |
| BT1496 | 95.977 | Dimethyl Sulfoxide | – – | 1 |
| BT3768 | 82.609 | Rhamnose monohydrate | – – – | 1 |

**B**

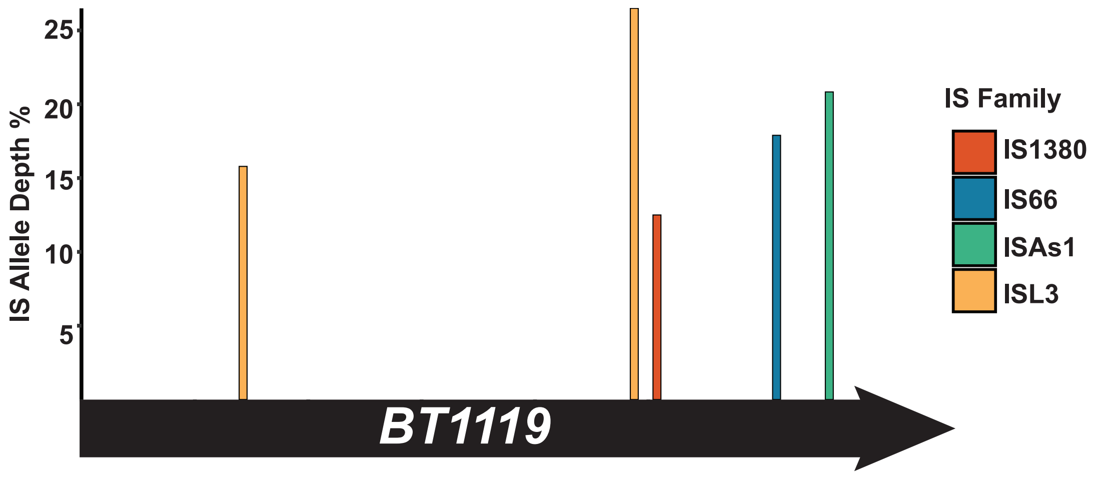


**Figure S3. IS element insertions in genes that affect Bt fitness, related to Figure 3. (A)** Table of IS element hits in genes involved in fitness in Bt. Genes whose fitness values < 0 are beneficial for fitness in the specified condition and genes whose fitness > 0 are detrimental to fitness in the specified condition (+++ fitness > 4, ++ fitness > 2, + fitness > 1, - - - fitness < -4, - - fitness < -2, - fitness < -1). **(B)** Schematic of IS element insertions in BT1119.

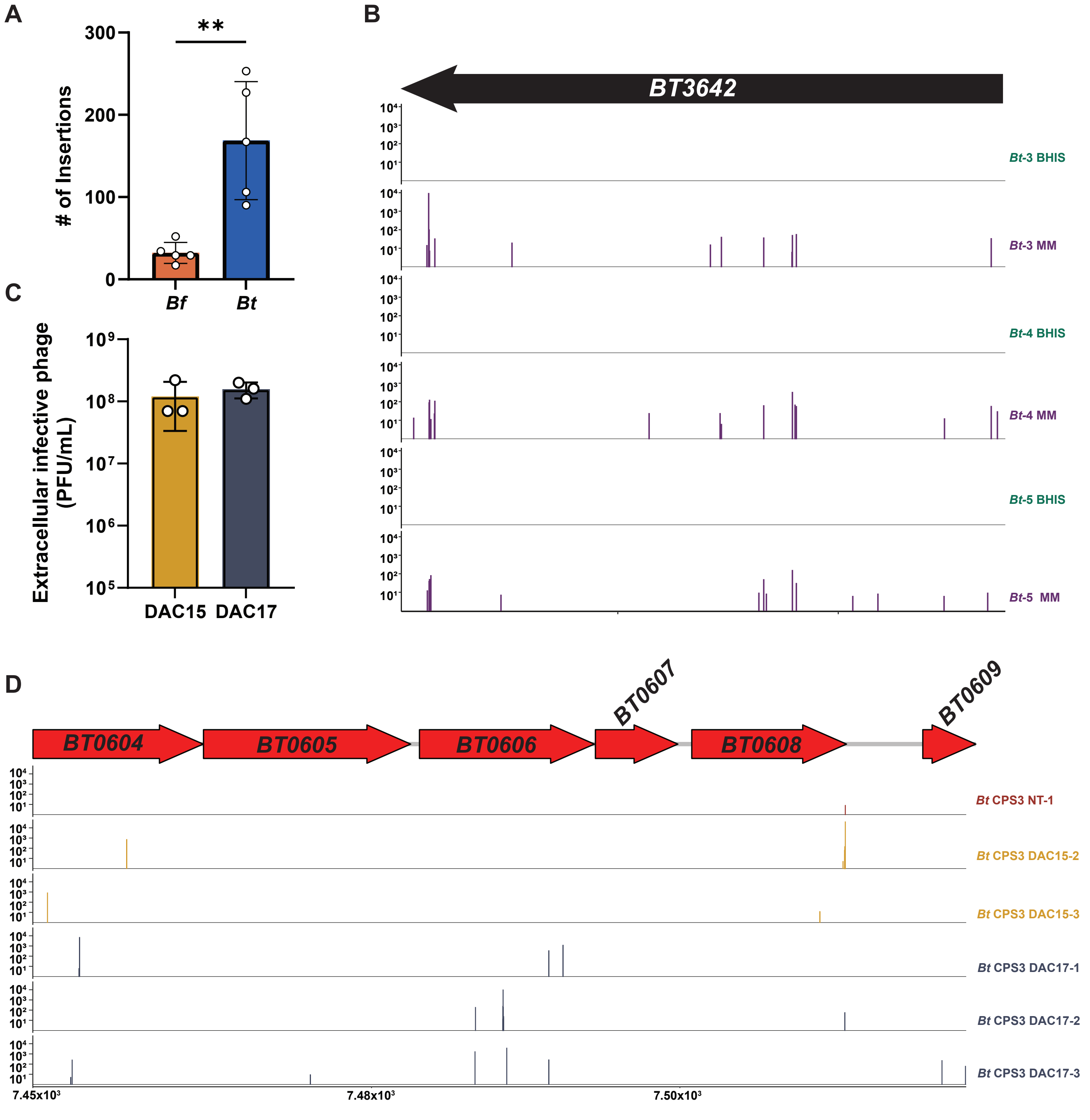


**Figure S4. ISOSDB412 drives IS element insertions during selective pressure, related to Figure 4. (A)** IS element insertions in Bt and Bf strains following passage in nutrient limited conditions (unpaired T test, **p < 0.01) **(B)** Schematic of IS element insertions in BT3642. **(C)** Quantification of infectious phage particles in the supernatant of Bt CPS3 strains chronically infected with DAC15 or DAC17. **(D)** Schematic of IS element insertions in the CPS3 locus.

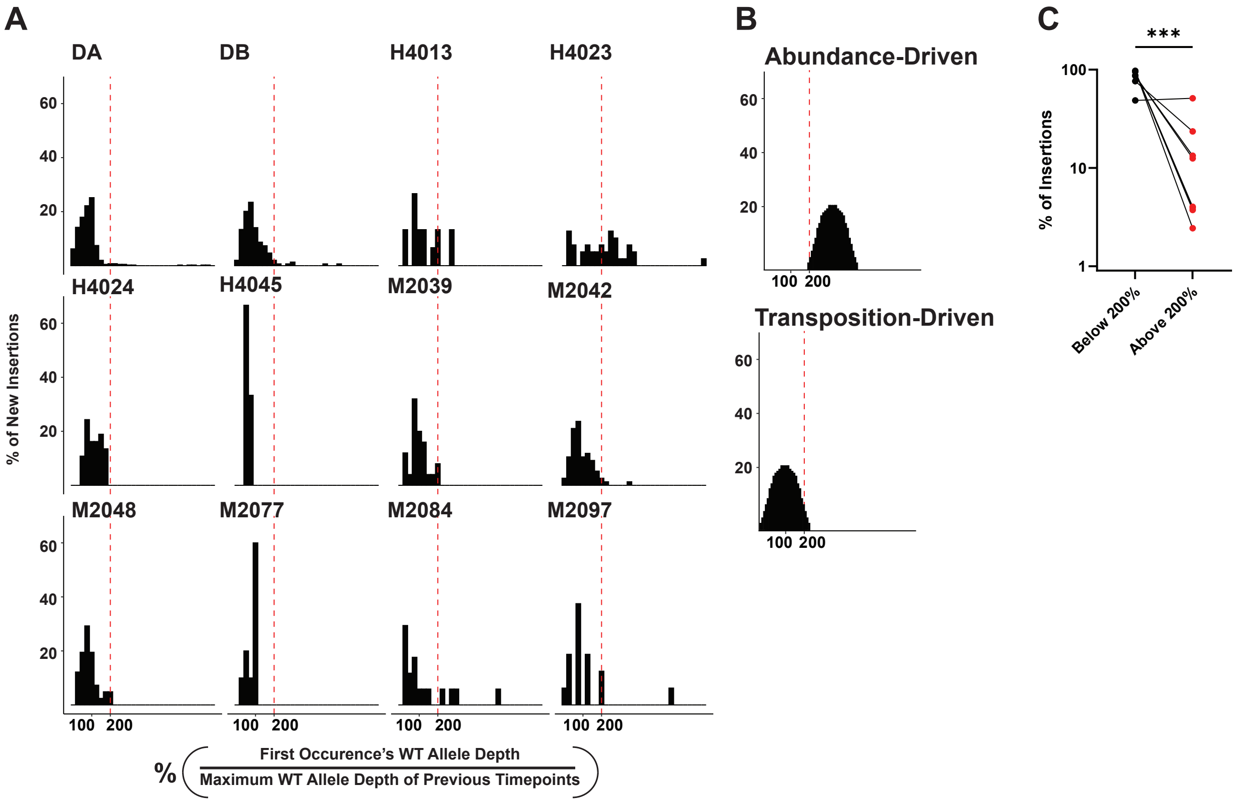


**Figure S5. New IS element insertions are not associated with increased wild type allele depth, related to Figure 5. (A)** The percentage of new IS element insertions is shown on the Y axis and the X axis is the percentage of the wild type allele depth from the first detection of each new insertion divided by the maximum detected wild type allele depth of the timepoints prior to the first detection timepoint. The red dashed line intersects the X axis at 200%. **(B)** Representation of abundance-driven or transposition-driven allele measurements. **(C)** Significantly more IS element insertions are predicted to arise from transposition compared to changes in host abundance (***$p < 0.01$, paired T-test).

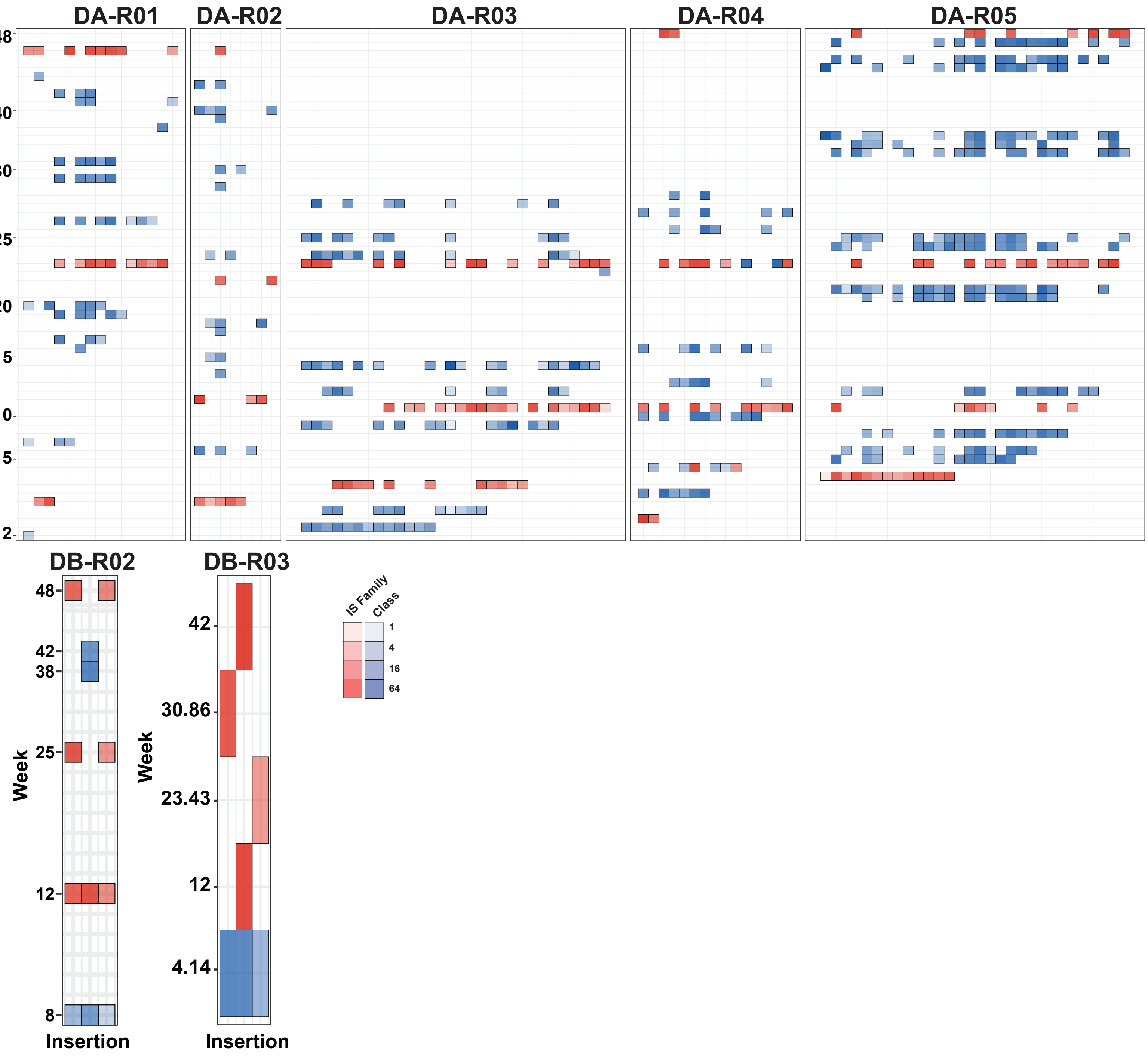


**Figure S6. Remainder of shared insertion plots, related to Figure 6.** Same scale in Fig. 6A is used in these figures.
